# MLL1 minimal catalytic complex is a dynamic conformational ensemble susceptible to pharmacological allosteric disruption

**DOI:** 10.1101/308676

**Authors:** Lilia Kaustov, Alexander Lemak, Hong Wu, Marco Faini, Scott Houliston, Lixin Fan, Xianyang Fang, Hong Zeng, Shili Duan, Abdellah Allali-Hassani, Masoud Vedadi, Ruedi Aebersold, Yunxing Wang, Cheryl H. Arrowsmith

## Abstract

Histone H3K4 methylation is an epigenetic mark associated with actively transcribed genes. This modification is catalyzed by the mixed lineage leukaemia (MLL) family of histone methyltransferases including MLL1, MLL2, MLL3, MLL4, SET1A and SET1B. Catalytic activity of MLL proteins is dependent on interactions with additional conserved proteins but the structural basis for subunit assembly and the mechanism of regulation is not well understood. We used a hybrid methods approach to study the assembly and biochemical function of the minimally active MLL1 complex (MLL1, WDR5 and RbBP5). A combination of small angle X-ray scattering (SAXS), cross-linking mass spectrometry (XL-MS), NMR spectroscopy, and computational modeling were used to generate a dynamic ensemble model in which subunits are assembled via multiple weak interaction sites. We identified a new interaction site between the MLL1 SET domain and the WD40 repeat domain of RbBP5, and demonstrate the susceptibility of the catalytic function of the complex to disruption of individual interaction sites.

## INTRODUCTION

Post-translational modifications on histone tails are key epigenetic signals for regulation of chromatin structure and gene expression. Histone H3 lysine 4 (H3K4) methylation is the epigenetic mark exclusively associated with transcriptionally active chromatin (1, 2). This modification is mostly catalyzed by the MLL/SET1 family of histone methyltransferases (3, 4), through their evolutionarily conserved SET domain (5, 6). The founding member of this family of H3K4 methyltransferases is the yeast SET1 protein (7, 8). In mammals, methylation of H3K4 is carried out by a family of six proteins: MLL (mixed lineage leukemia protein)1 to MLL4, SET1A and SET1B (9–15). The MLL proteins play crucial roles in embryonic development and hematopoiesis through transcriptional regulation of the clustered homeobox *(Hox)* genes and other genes important for developmental regulation (10, 16–19). Deletion of MLL1 and MLL2 can lead to severe defects in embryonic development in mice (18, 20). The *MLL1* gene is frequently rearranged in human acute leukemia in both adults and children (21–23). MLL3 and MLL4 have also been linked to other human malignancies. Recently studies have identified inactivating mutations in MLL3 and MLL4 in different types of human tumors (24–27), as well as in Kabuki syndrome (28).

The catalytic activity of MLL family members are dependent to varying degrees on the presence of additional conserved protein subunits, RbBP5, WDR5 and ASH2L, and a minimal core enzyme can be reconstituted with the conserved C-terminal SET domain of MLLs and at least two of the other subunits. Interestingly, in studies of these reconstituted core enzymes, MLL1 appears to be unique among the family members in its requirements for, and interactions with other subunits. For example, compared to other family members, the catalytic activity of MLL1 is the most dependent on WDR5 (29–31). Similarly, MLL1 binds with the least affinity to the RbBP5-Ash2L heterodimer, and its catalytic activity is only weakly stimulated by RbBP5-ASH2L compared to WDR5 (32).

Recent crystallographic studies of MLL3 support a model in which the RbBP5-ASH2L heterodimer stabilizes the catalytically active conformations of MLL2,3,4 through interactions with conserved surfaces on their SET domain (32). However, it was suggested that two key variant residues on this surface of MLL1 dramatically weakened the interaction between MLL1 and RbBP5-ASH2L relative to that of other MLL members, thereby increasing the dependence of MLL1 on the WDR5 subunit. The unique dependence of MLL1 activity on WDR5 may be of therapeutic relevance, as we and others have shown that pharmacological targeting of the MLL interaction site on WDR5 can functionally antagonize MLL1 in cancers that are dependent on MLL1 activity (33–35). While there are several structures of WDR5 bound to MLL and RbBP5 peptides (31, 36–39), as well as a crystal structure of the apo-SET domain (40) of MLL1 and a 24 Å resolution cryo-EM model of the homologous yeast COMPASS complex (41), an atomic level picture of a functional MLL1 catalytic core complex is still lacking. Here, we report a hybrid methods study of MLL1 and its catalytic core components in solution. Using small angle X-ray scattering (SAXS), cross-linking mass spectrometry (XL-MS), NMR spectroscopy, and computational modeling we derived a dynamic ensemble model for MLL1/WDR5/RbBP5 and identify a new interaction site between MLL1-SET domain and the N-terminal WD40 repeat domain of RbBP5. Our data support the notion that the functional MLL1 enzyme comprises a collection of weak but specific interactions, and that the disruption of individual interactions can have significant destabilizing effects on the entire complex. These results highlight the dynamic nature of an important protein complex and the strategy of targeting a weak but druggable protein-protein interaction site to antagonize the function of a larger macromolecular assembly that is dependent on a collection of weak interactions.

## MATERIALS AND METHODS

### Cloning of MLL1, WDR5 and RbBP5 constructs

The coding regions for human MLL1-SET (aa3785-3969) and MLL1-WIN (aa3745-3969) were PCR-amplified and sub-cloned into the pET28GST-LIC vector (GeneBank ID: EF456739). The mutants, MLL1-SET-7D (Δ3786-3793) and MLL1-SET _Q3787V/P3788L/-Y3791G, were generated from the wild-type clone using the QuikChange PCR mutagenesis kit (Agilent). RbBP5 constructs of different lengths (aa 10-340, 320-410, 340-538, 1-538) and WDR5 (aa 24-334) were sub-cloned into the pET28-MHL vector (GeneBank ID: EF456738). For expression of dimeric complexes and reconstitution of the trimeric MLL1 complex, full-length WDR5 and MLL1-WIN, WDR5 and RbBP5 were cloned into pFastBac Dual expression vector (Thermo Fisher Scientific), respectively, with MLL1-WIN and RbBP5 tagged with an N-terminal hexa His-tag.

### Protein preparation

The individual components of the MLL1 complex used in this study were expressed in *E. coli* and purified using an N-terminal GST-tag (for MLL1) or His-tag (for WDR5 and RbBP5). The dimeric and trimeric complexes of MLL1 used for SAXS and cross-linking studies were expressed in Sf9 cells. The dimeric complex of WDR5-MLL1-WIN and WDR5-RbBP5 were purified using TALON affinity resin (Clontech) followed by gel filtration chromatography. The purified dimeric complexes were incubated on ice for 2 hours together to reconstitute the trimeric complex which was purified and recovered by gel filtration chromatography. Detailed procedures are described in the Supplementary Data section.

### SAXS data collection, analysis and modelling

SAXS measurements were carried out at the beamline 12-ID-C of the Advanced Photon Source, Argonne National Laboratory. The energy of the X-ray beam was 18 Kev (wavelength λ=0.6888 Å), and two setups (small– and wide-angle X-ray scattering, SAXS and WAXS) were used in which the sample to charge-coupled device (CCD) detector (MAR research, Hamburg) distance were adjusted to achieve scattering *q* values of 0.006 < *q* < 2.3Å^-1^, where q = (4π/λ)sinθ, and 2θ is the scattering angle. Data were analyzed using the program PRIMUS (ATSAS package, EMBL (42)). Detailed descriptions of SAXS data collection and analysis, and modelling protocols, are provided in the Supplementary Data.

### Chemical cross-linking mass spectrometry

The reconstituted trimer complex of WDR5, RbBP5 and MLL1-SET was cross-linked at a concentration between 12 and 16 μM, with 1 mM of isotopically coded disuccinimidyl suberate (DSS-d_0_,DSS-d_12_) as described previously (43). Protease digestion was carried out with LysC and trypsin. After acidification, cross-linked peptides were purified on C18 cartridges and enriched by size-exclusion chromatography (SEC). SEC fractions were analyzed in duplicate on an LC-MS (Easy-nLC 300; Orbitrap LTQ XL). For complete details refer to Supplementary Data.

### NMR Spectroscopy

NMR spectra were collected at 25°C on a Bruker Avance(II) 800 MHz spectrometer equipped with a cryoprobe. Samples contained 5% D_2_O with protein concentrations ranging from 100 to 350 μM. For the assignment of the backbone resonances of the ΔN-WDR5 construct, a triply-labeled (^15^N/^13^C/^2^H) sample was prepared and conventional triple-resonance backbone spectra were acquired as described previously (44), in order assign backbone and Cβ resonances using the ABACUS approach (45). [^1^H-^15^N]-TROSY titrations of ^15^N-labeled WDR5 was performed by adding aliquots of MLL1-WIN (GSARAEVHLRKS) and RbBP5-WBM (EDEEVDVTSV) peptides at molar ratios ranging from 1:1 to 1:7. The weighted chemical shift displacements were calculated using the following formula: Δppm=[δ_NH_)^2^+(δ_N_/5)^2^]^1/2^. Spectra were processed with NMR Pipe (46) and analyzed with SPARKY (47).

### GST Pull-down experiments

Recombinant purified MLL1-GST proteins were incubated with RbBP5 constructs in an assay buffer containing 20mM TRIS pH 7.7, 150 mM NaCl, 10μM ZnCl_2_, 5mM β-mercaptoethanol, 5 mM DTT, 1 mM PMSF in a 1:2 molar ratio at 4°C for 1h. Proteins were then incubated with 100μl of glutathione-Sepharose beads (GE Healthcare) for an additional 1h. The mixture was transferred to a micro-column and was extensively washed with assay buffer. Bound proteins were eluted with 30mM reduced glutathione and detected by SDS-PAGE and Coomassie staining.

### Histone methyltransferase assay

Activity assays were performed in 50 mM Tris-HCl, pH 8.0, 5 mM DTT and 0.01% Triton X-100, using 5 μM ^3^H-SAM and 5 μM Biotin-H3 1-25. Increasing concentrations of RbBP5 were added to 200 nM of MLL1-WDR5 complexes (with either wild-type or mutant MLL1). All reactions were incubated for 90 minutes at room temperature and the SPA method was used to determine activities. Experiments were performed in triplicate. To test the effect of OICR-9429 on the MLL1 complex, increasing concentrations of the compound was incubated with 200 nM MLL1-WDR5 complex for 20 min before adding 400 nM RbBP5. The activity of the complex was measured as above.

## RESULTS AND DISCUSSION

### SAXS data reveal solution ensembles for WDR5, RbBP5 and MLL-SET

To model catalytically active MLL1 complexes, we first collected reference solution data for the individual subunits including the SET domain of MLL1, the WD40 repeat region of WDR5 (ΔN-WDR5), the N-terminus of RbBP5 (RbBP5-NTD) and full-length RbBP5, followed by characterization of dimeric and trimeric complexes. Fig 1A shows the protein constructs used in this study. Normalized Kratky plots of ΔN-WDR5 and RbBP5-NTD exhibit a typical bell-shape expected for a globular protein and are nearly superimposable in the q range 0<qRg<3 (Fig 1B). Also, the experimental values of Rg predicted for ΔN-WDR5 and RbBP5-NTD are in agreement with the theoretical values expected for globular proteins (Table 1 and Fig S2). The normalized Kratky plot of MLL1-SET also exhibits a bell-shape, but its maximum is shifted with respect to the globular protein position, with poor convergence at high q-values, indicating that MLL1-SET is flexible. The observed flexibility of the MLL1-SET could be attributed to known inherent dynamics of the SET domain in the absence of cofactor (32), and to the disordered N-terminal tail. The calculated solution ensembles for each protein taking into account known or predicted disordered regions (see SI for details) establish good correspondence between our SAXS measurements and the crystal structures of WDR5, the SET domain of MLL1, and our homology model of RbBP5-NTD predicted using ROSETTA (48)(Fig S1).

**Figure 1.**
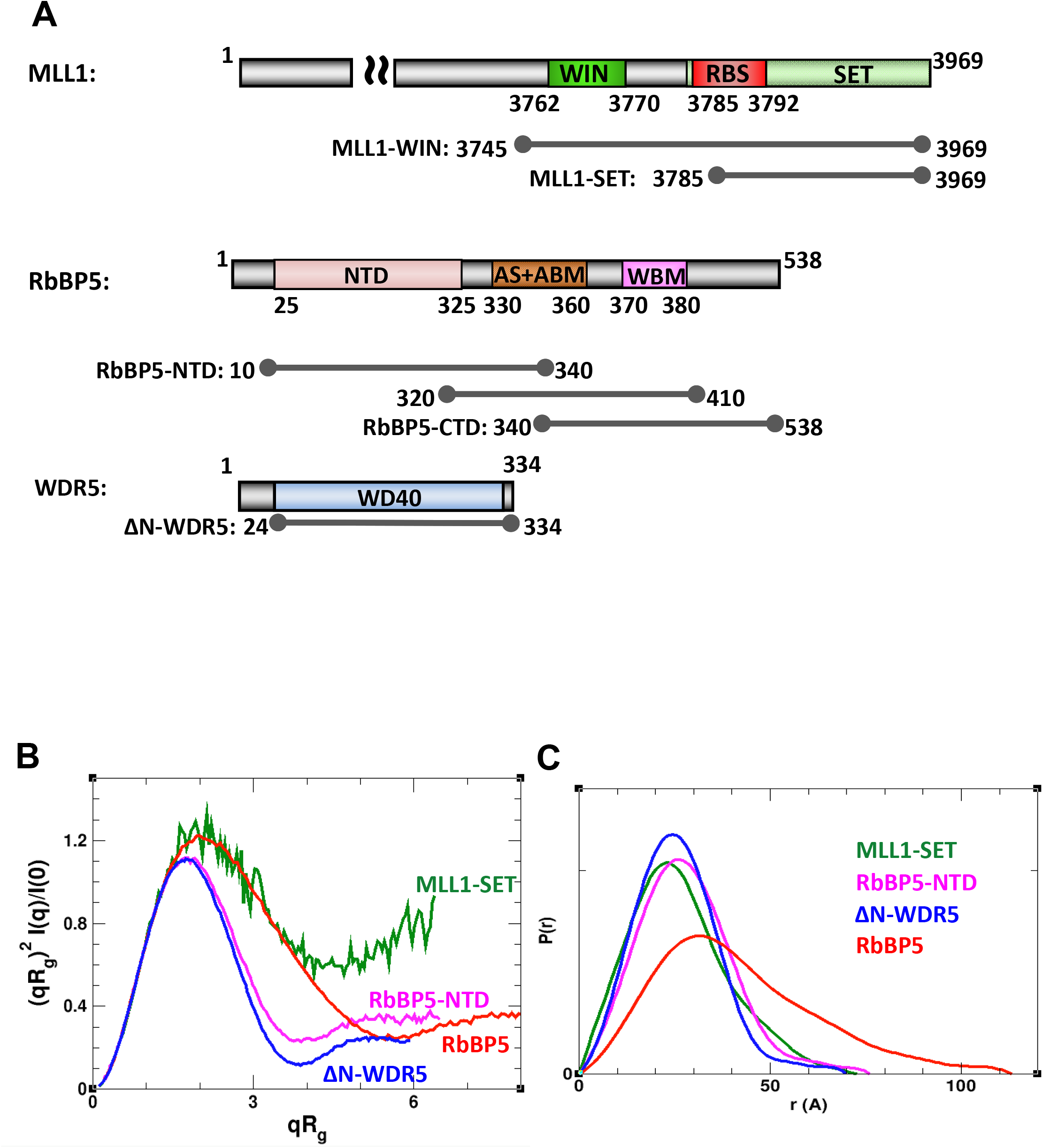
Individual components of the WDR5-RbBP5-MLL1 core complex. (A) Schematic representation of MLL1, RbBP5 and WDR5 domain organization and constructs used in this study. For clarity, only the C-terminal region is shown for MLL1. WIN: WDR5 interacting motif as previously defined (36); RBS: RbBP5 binding site as defined in this study; SET: catalytic methyltransferase domain; NTD: N-terminal domain; AS+ABM: activation segment and ASH2L binding motif as defined in (32); WBM: WDR5 binding motif (38). (B) Rg-based Kratky plots of experimental SAXS data for MLL1-SET (green), ΔN-WDR5 (blue), RbBP5-NTD (magenta), and full-length RbBP5 (red). (C) Normalized pair distance distribution function P(r) calculated from experimental SAXS data with GNOM. See also Table 1 and Figs S1, S2.

**Table 1.** SAXS parameters for data validation and interpretation.

|  | **Rg<sup>a</sup>** (Å) | **Rg<sup>b</sup>** (Å)<br>(real) | **D<sub>max</sub><sup>c</sup>** (Å) | **V<sub>c</sub><sup>d</sup>** | **Mw<sup>e</sup>** (kDa) | **NSD<sup>f</sup>** (SAXS<br>envelope) |
| --- | --- | --- | --- | --- | --- | --- |
| *Individual* |  |  |  |  |  |  |
| **MLL1-SET<sup>g</sup>** | 20.2 | 20.8 | 73 | 207 | 22.0 (21.6) | 0.58 |
| **ΔN-WDR5<sup>h</sup>** | 19.8 | 19.9 | 70 | 251 | 25.9 (34.1) | 0.60 |
| **RbBP5-NTD<sup>i</sup>** | 21.5 | 21.7 | 76 | 276 | 28.7 (36.8) | 0.72 |
| **RbBP5<sup>j</sup>** | 32.1 | 33.0 | 113 | 489 | 60.6 (59.1) | 0.64 |
| *Dimeric complexes* |  |  |  |  |  |  |
| **WDR5/**<br>**MLL1-WIN<sup>k</sup>** | 32.5 | 33.6 | 120 | 372 | 56.5 (64.7) | 0.79 |
| **WDR5/RbBP5<sup>l</sup>** | 39.8 | 41.1 | 140 | 669 | 91.2 (96.5) | 0.79 |
| *Trimeric complex* |  |  |  |  |  |  |
| **WDR5/RbBP5/**<br>**MLL1-WIN<sup>m</sup>** | 49.1 | 51.8 | 183 | 929 | 135.5 (124.6) | 0.69 |
| Oi |  |  |  |  |  |  |

a) Radius of gyration calculated using Guinier fit
b) Radius of gyration calculated using GNOM (51)
c) Maximum distance between atoms from GNOM
d) Volume of correlation (52)
e) Molecular weight estimated from SAXS data using V<sub>c</sub> (52). The MW expected from the sequence is shown in the parentheses
f) NSD: Normalized spatial discrepancy; The values given are the average and standard deviation from fifteen runs of DAMMIF (53)
g) MLL1<sub>3785-3969</sub>
h) WDR5<sub>524-334</sub>
i) RbBP5<sub>10-340</sub>
j) RbBP5 (full-length)
k) full-length WDR5 and MLL1<sub>3745-3969</sub>
l) full-length WDR5 and RbBP5
m) full-length WDR5, full-length RbBP5, and MLL1<sub>3745-3969</sub>

One of the main challenges in modeling the MLL1 complex is the lack of structural information on RbBP5. To better understand its structural arrangement, we collected [^1^H-^15^N]-TROSY NMR spectra of a full-length construct, as well as constructs corresponding to its C-terminal and N-terminal regions (Fig 2A). The data confirm that RbBP5-NTD is a globular, folded domain, consistent with our SAXS analysis and WD40 homology model. The C-terminal region of RbBP5 (RbBP5-CTD) is substantially disordered as evidenced by the lack of spectral dispersion (Fig 2A). Both the gel filtration profile (Fig S3B) and the radius of gyration estimated from SAXS data (Table 1 **and** Fig S2) indicate a high degree of disorder in the RbBP5 full-length protein. This is further supported by sequence based secondary structure prediction and order parameters, which predict a rigid globular N-terminus and a flexible coil-like C-terminus (Fig S3). Interestingly, the [^l·^H-^15^N]-TROSY spectrum of full-length RbBP5 is not the superposition of the individual NTD and CTD spectra, and reflects features of both folded and unfolded regions with some apparent conformational broadening, possibly reflecting weak intramolecular interactions (Fig S3A).

**Figure 2.**
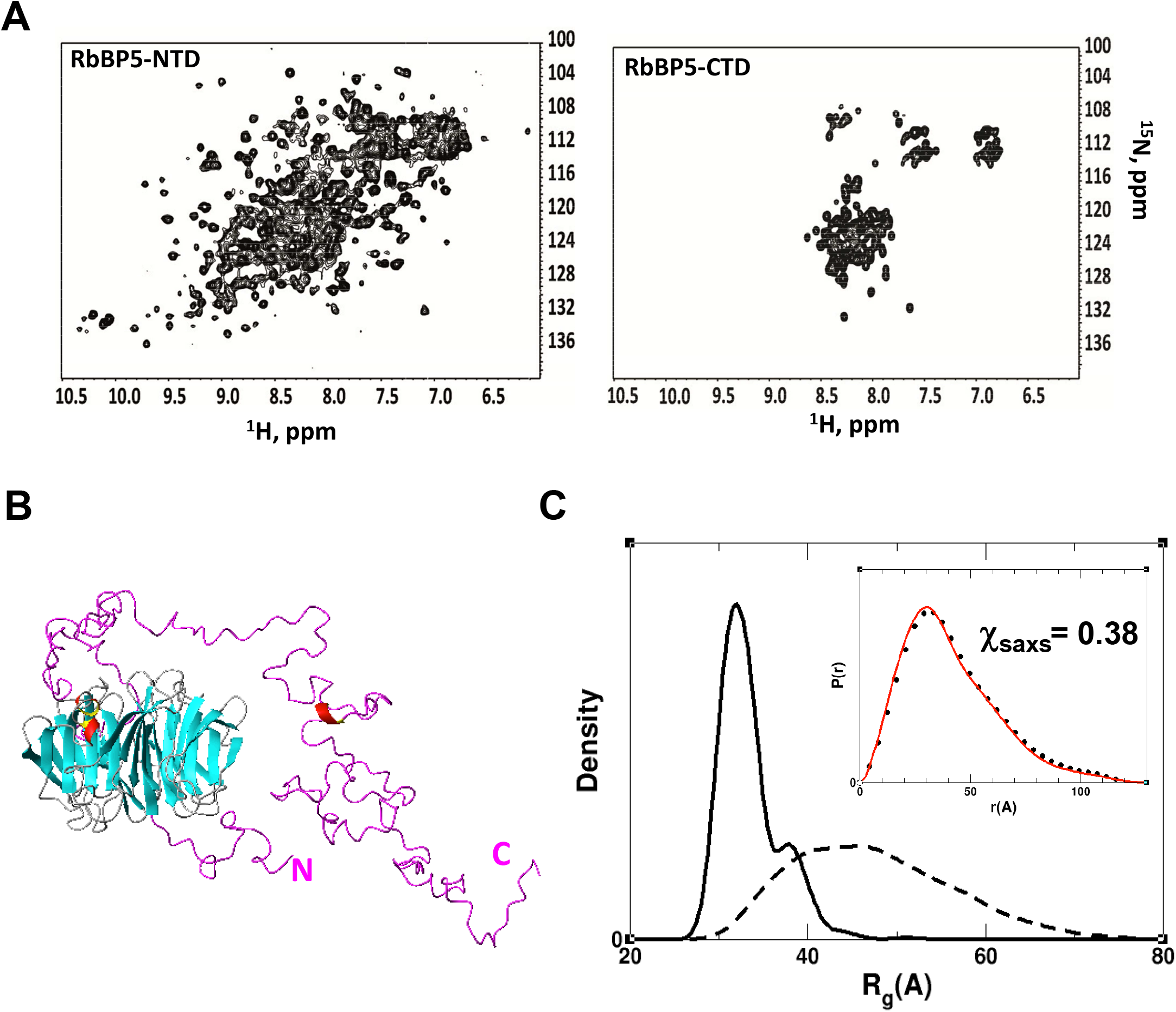
SAXS model for RbBP5 full-length protein. (A) [^1^H-^15^N]-TROSY spectra of RbBP5-NTD (left) and RbBP5-CTD (right) constructs. For the CTD, amide resonances are clustered between 7.5 and 8.5 ppm in the ^1^H dimension indicating that it is unfolded, while spectra for the NTD is consistent with a structured domain. (B) A representative member of the most populated SES ensemble model reflects a structured N-terminal domain and a flexible, but non-random, C-terminus. (C) The clear difference in Rg distribution profiles for the initial pool of 30,000 models (with random conformations of the C-terminal part (dashed)) *vs*. the SAXS-derived SES ensemble (solid) indicate that the CTD, in the context of full-length RbBP5, is not randomly disordered. Pair distance P(r) distribution function calculated for the experimental data (black circles) and for the SAXS-derived SES ensemble (red line) are displayed in the inset. See also Figs 1C, S1–S3.

The NMR data is consistent with the general shape of the normalized Rg-based Kratky plot and the pair distance distribution function P(r) for full-length RbBP5 (Fig 1B, C). In particular, the P(r) function of RbBP5 has an asymmetric shape with a long smooth tail at large r values, and the position of its maximum is shifted only slightly (~4 Å) with respect to that of RbBP5-NTD. The latter features indicate that full-length RbBP5 has no additional globular content compared to RbBP5-NTD.

Based on the above data we used the Sparse Ensemble Selection (SES) approach (49) to calculate a solution ensemble of RbBP5 that would satisfy the SAXS data. An initial ensemble consisting of 20,000 models of RbBP5 with random conformations of its flexible regions (residues 1-23 and 326-538) did not fit the SAXS data well (the goodness-of-fit χ=9.4). We next generated a solution ensemble that better fits the SAXS data by calculating an optimal weight for each model in the initial ensemble using a multi-orthogonal matching pursuit algorithm (49) (see SI for details). The resulting optimal ensemble fits the SAXS data very well with χ=0.38 (Figs 2C, S1). The most populated models in the optimal ensemble are shown in Figs 2B, S1F, G. The optimal ensemble displays a much narrower distribution of radius of gyration values than the initial random ensemble, with a major peak at Rg=37 Å (Fig 2C). This indicates that RbBP5 adopts a more compact conformation than would be predicted for a fully random CTD, consistent with our NMR data for the full-length protein.

### Binary subcomplexes have dynamic non-random solution conformations mediated by WD40 repeat domains

Our SAXS data for the binary complexes of WDR5/MLL1-WIN and WDR5/RbBP5 both suggest the presence of significant disorder, especially for WDR5/MLL1-WIN (Fig 3A, (S3A). The P(r) functions of WDR5/MLL1-WIN and WDR5/RbBP5 are typical for proteins containing globular domains tethered by long disordered regions (Fig 3A). The position of the P(r) major peak for the aforementioned complexes is close to the positions of the major peaks of P(r) of their individual components (Fig 1C), indicating that in both complexes the globular domains are not in close contact and may not adopt a unique arrangement in solution. WDR5 is known to interact with RbBP5 and MLL1 through small peptide segments designated as the WDR5 binding motif (WBM) (38) and WDR5 interacting (WIN) (36) motif, respectively (Fig 1A). Both interactions with WDR5 have reported dissociation constants on the order of 1-2 μM (30, 36, 38, 39). To calculate solution ensembles of the WDR5 binary complexes, we first used NMR to verify that WDR5’s mode of interaction with these two motifs, as observed in the crystal structures, is maintained in solution. We expressed a triply-labeled (^15^N/^13^C/^2^H) ΔN-WDR5 construct, and were able to assign 254 amides (Fig S6). We then used chemical shift perturbation (CSP) analysis in [^1^H-^15^N]-TROSY titration experiments, to localize the WRD5 binding site for peptides corresponding to the two motifs. As seen in Fig 3B, there is excellent agreement between the WDR5 CSP profiles and the WDR5/WIN (PDB:4ESG) and WDR5/RbBP5 (PDB:2XL2) crystal structures. Next, using these two structures to fix each WDR5-peptide interface, we modeled the ensemble of solution conformations for the binary complexes of WDR5 with MLL1-WIN and full-length RbBP5 using the SES method. The arrangement of the globular domains in the most populated models of the optimal ensembles for both WDR5/MLL1-WIN and WDR5/RbBP5 complexes does not support the existence of additional interactions of WDR5 with MLL1-WIN or with RbBP5 other than those described above (Fig S4).

**Figure 3.**
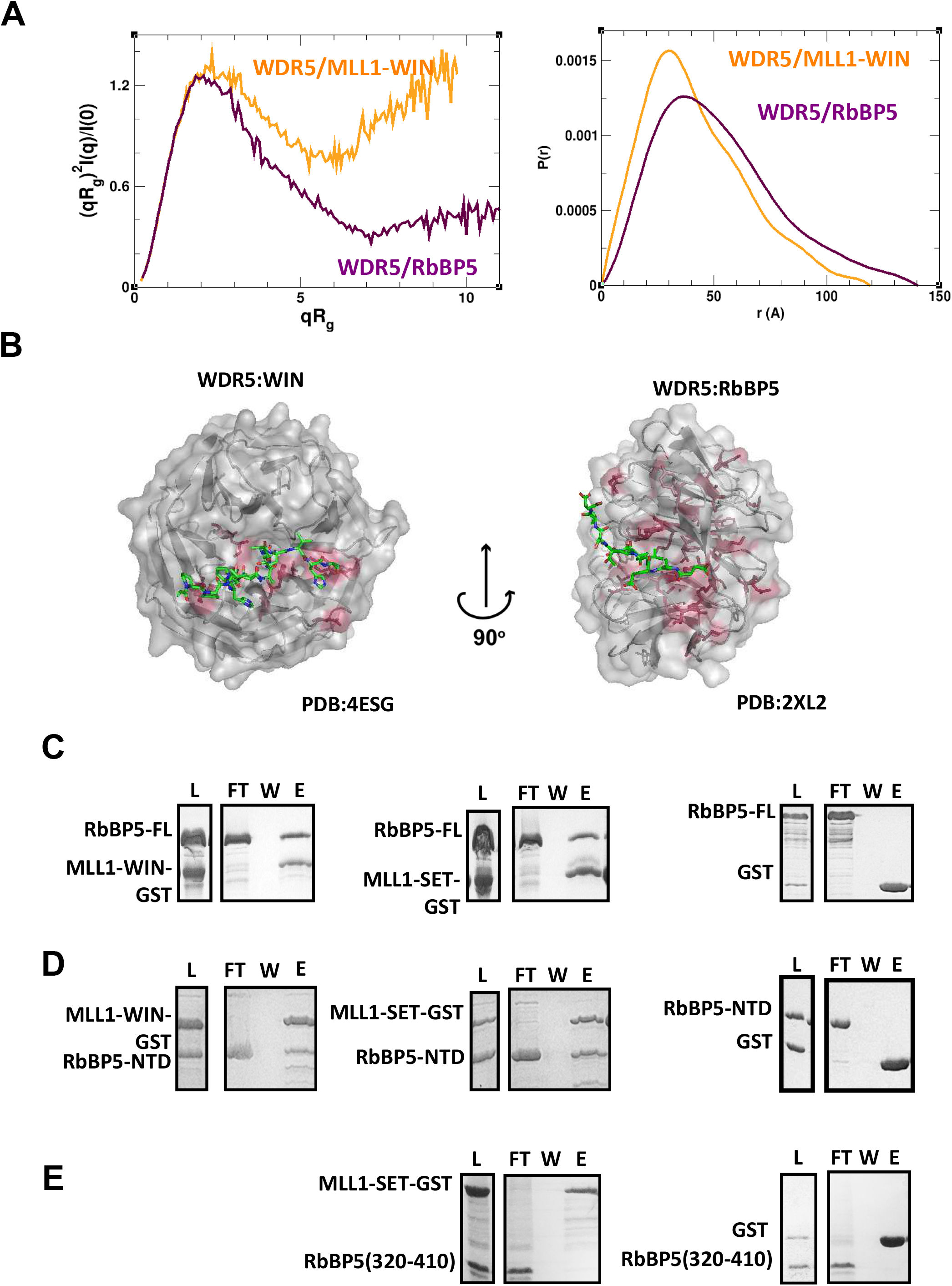
WDR5/MLL1 and RbBP5/MLL1 Binary Complexes. (A) Rg-based Kratky plots of experimental SAXS data for binary complexes of WDR5/MLL1-WIN (orange) and WDR5/RbBP5 (maroon). (B) WDR5 residues (pink) that show substantial CSPs in [^1^H-^15^N]-TROSY spectra when titrated with WIN and RbBP5 peptides (green). The perturbation patterns were mapped onto, and are in excellent agreement the WDR5/WIN (PDB: 4ESG) and WDR5/RbBP5 (PDB:2XL2) structures. (C-E) Interaction between RbBP5 constructs and MLL1-WIN/MLL1-SET. GST-pull down experiments show direct interaction between (C) RbBP5-FL and (D) RbBP5-NTD with MLL1 constructs along with the control interaction between (C) RbBP5-FL and (D) RbBP5-NTD with MLL1 constructs along with the control GST experiments. (E) There was no interaction detected between RbBP5_320-410_ and MLL1-SET. Lanes correspond to L=loaded protein mixture; FT=flow through; W=wash; E=eluate. See also Figs S3, S4.

There is currently no atomic resolution structural data for the interaction of MLL1 with RbBP5. Recently, the activation segment (AS) of RbBP5 was shown to bind to the SET domain of other MLL family members, but only very weakly to MLL1 (32). In order to determine whether there is a direct interaction between RbBP5 and MLL1, we performed GST pull-down studies of full-length RbBP5, RbBP5-NTD and RbBP5 (320-410) with GST-MLL1-SET and GST-MLL1-WIN (Fig 3C-E). Both MLL1 constructs interacted with RbBP5 constructs containing the N-terminal WD40 repeat but did not interact with the C-terminal residues (aa 320-410) of RbBP5 containing the AS region. These results agree with the lack of conservation of the RbBP5 AS-binding surface on MLL1 (32), and suggest that MLL1-SET may interact with the WD40 domain of RbBP5.

### SAXS and cross-linking data suggest a dynamic triangulated ensemble for WDR5/RbBP5/MLL 1 – WIN

Our SAXS data for the catalytically active WDR5/RbBP5/MLL1-WIN complex showed a substantial amount of flexibility. The shape of the experimental Kratky plots of the complex is typical of partially disordered proteins (Figs 4A, S5A). In particular, the Rg-based Kratky plot is a bell-shaped curve with a maximum at (2.26, 1.27) shifted to higher values of the coordinates with respect to its position expected for a globular protein. Also, the presence of a high degree of flexibility is evidenced by the poor convergence of the Kratky plots at high q values. The low maximum value of 0.48 in the Vc-based Kratky plot (Fig S5A), as well as the asymmetric shape of the P(r) function (Fig S5B), suggests an elongated shape of the complex. This is in agreement with the averaged *ab initio* SAXS-derived molecular envelope, which showed an extended shape with approximate dimensions of 220×105×70 Å (Fig 4D).

**Figure 4.**
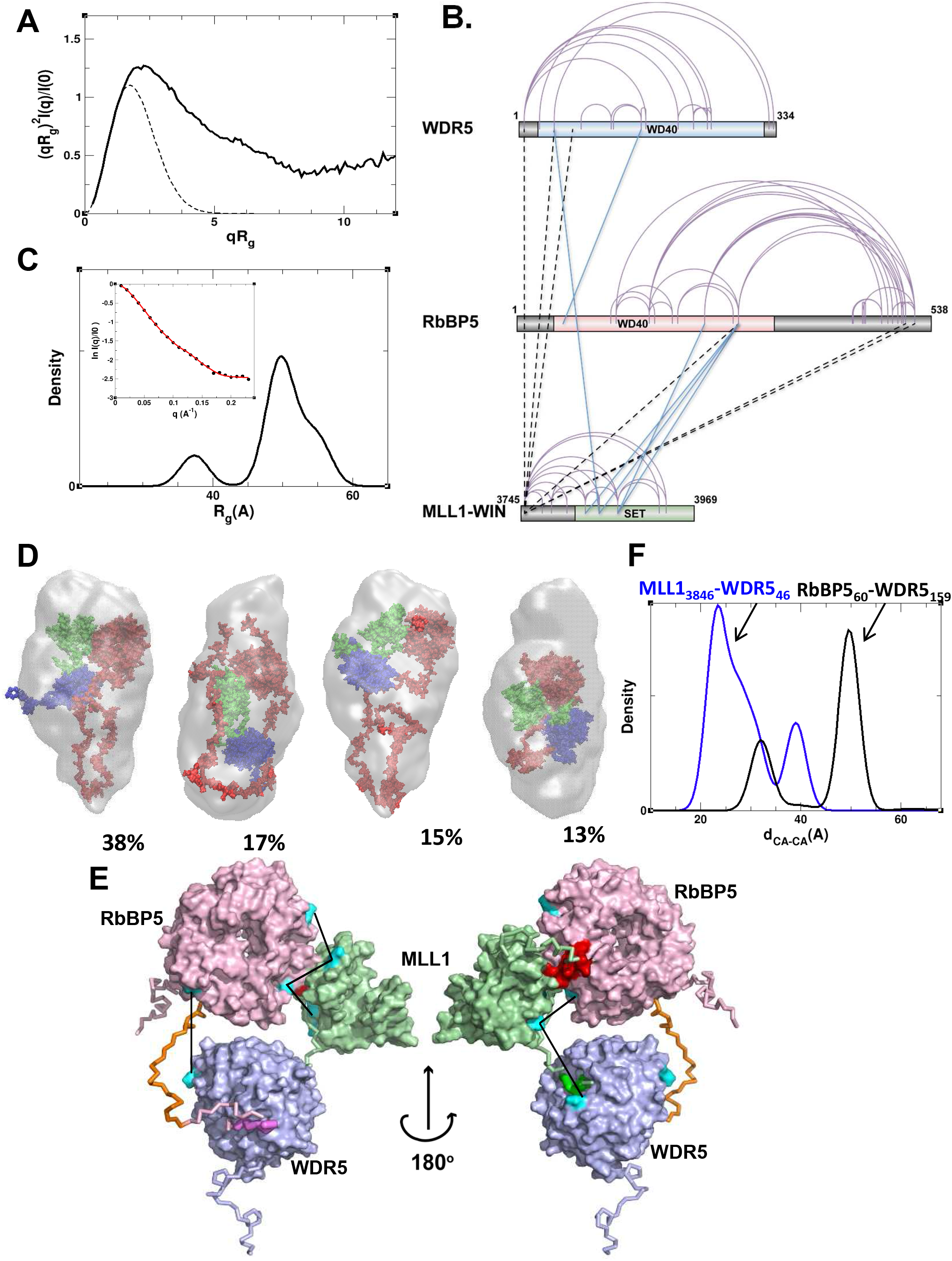
Dynamic model of trimeric WDR5/RbBP5/MLL1-WIN complex derived from SAXS and cross-link data. (A) Rg-based Kratky plot of experimental SAXS data for WDR5/RbBP5/MLL1-WIN indicates a high degree of flexibility within the complex. (B) Sequence mapping of intra-protein and interprotein cross-links. Intra-protein cross-links are indicated with purple arcs. Inter-protein crosslinks between globular domains are indicated with green lines and inter-protein cross-links within flexible regions are indicated with black dashed lines. (C) Rg distribution for the optimal ensemble of trimeric complex models is shown by a solid black line. Experimental SAXS profiles (black circles) plotted with the theoretical profiles (red line) averaged over the ensemble are shown in the inset. (D) Surface representation of the four most populated models in the optimal ensemble models of WDR5/RbBP5/MLL1-WIN overlaid with average *ab-initio* SAXS-predicted molecular envelope (gray mesh). WDR5, MLL1 and RbBP5 are colored in blue, green and red, respectively. (E) The most populated (~40%) model of the complex is shown by a surface representation of the globular regions and a backbone trace of the flexible regions. For clarity, the C-terminus of RbBP5 (i.e. RbBP5_382-538_) is not shown and the residues from the AS+ABM region are in dark orange, WIN in green, WBM in violet and RBS in red. The crosslinks between globular domains are shown by solid black lines and the cross-linked lysine residues are shown in cyan. (F) The ensemble distribution of the distance between C_α_ atoms of Lys residues involved in cross-links between MLL1-SET and WDR5, RbBP5-NTD and WDR5 are shown by solid blue and black lines, respectively. See also Fig S5.

We note that pair-distance distribution functions of proteins containing several globular domains connected by long disordered regions are characterized by peaks at low r-values, corresponding to the intra-domain distances. Therefore, if the three globular domains of WDR5, MLL1-SET and RbBP5-NTD are not interacting directly with each other within the trimeric MLL1 complex, we would expect the P(r) function of the complex to have peaks at 26-32 Å, reflecting the inter-atomic distances prevailing within these domains (Figs 1C, (S5B). However, the experimental P(r) function of the trimeric MLL1 complex has its maximum at a much larger distance of ~ 47 Å (Fig S5B), suggesting the existence of possible inter-domain contacts in the complex.

To aid our modeling of the solution conformations of the trimeric complex we performed cross-linking mass spectrometry studies. We observed many intramolecular cross-links within each of the three proteins. These were highly consistent with the available WDR5 and MLL1-SET crystal structures, and importantly, in agreement with our RbBP5-NTD homology model, establish that these structural models are reliable representations of the domains within the trimeric complex in solution. We also observed a number of intermolecular cross-links, with the largest number being between MLL1 and RbBP5, suggesting association of these two subunits in solution. Fig 4B shows the sequence mapping of both intra– and inter-molecular DSS cross-links observed for the trimeric complex. For the purposes of modeling we used only intermolecular cross-links between lysine residues within the globular subunits (Table S1). These six cross-links are shown on Fig 4B by solid blue lines.

Using both SAXS and cross-linking data as conformational restraints we utilized the SES approach to calculate solution ensembles of the WDR5/RbBP5/MLL1-WIN complex that would satisfy both sets of experimental data. An initial pool of representative structures was generated by combining rigid-body modeling and molecular dynamics simulations for both all-atomic and coarse-grained models of the complex. It was assumed that the MLL1-SET and RbBP5-NTD domains were tethered to WDR5 by WIN and WBS motifs, respectively, as seen in crystal structures (30, 36–38), along with cross-link derived distance restraints (see SI for details). It should be noted here that individual members of this initial ensemble did not necessary satify all inter-molecular cross-links: each satisfies on average 3 to 4 cross-links.

The optimal SES ensemble of WDR5/RbBP5/MLL1-WIN fits the SAXS data as a whole, with 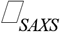 = 0.23 in the q-range 0<q<0.23. While only SAXS data were used to select the optimal ensemble, each inter-domain cross-link is consistent with at least one member of the ensemble, so that the ensemble as a whole is consistent with all cross-links. The most highly populated models of the optimal SES ensemble are shown in Figs 4 and S5. The optimal ensemble contains two clear populations; a smaller population (~13%) with a radius of gyration centered at ~37 Å and a more predominant population (~85%) with a more widely distributed radius of gyration centered at ~ 52 Å (Fig 4C, D). The relative position of the MLL1-SET and RbBP5-NTD domains is well defined by four inter-domain cross-links and their association is similar in all models within the optimal ensemble. In contrast, the relative position of WDR5 (compared to RbBP5-NTD and MLL1-SET) varies within the ensemble due to flexible linkers between the globular RbBP5-NTD and MLL-SET domains and their WDR5 interacting sequences (WBS and WIN, respectively) (Fig 4E, (S5C). There are only two inter-domain cross-links that involve WDR5, and they can only be simultaneously satisfied in the more compact subpopulation corresponding to Rg ~ 37 Å. The ensemble distribution of Cα-Cα distances corresponding to these cross-links showed a large fraction of ensemble members for which the cross-links cannot be simultaneously formed (Fig 4F).

### RbBP5-NTD has a unique interaction mode with MLL1

A recent crystallographic study revealed an important role for the AS+ABM region of RbBP5 in binding to the SET domain of MLL family proteins, thereby stimulating the latter’s catalytic activity (32). This work showed that the catalytic activity of MLL2, 3, 4 and SET1A/B was highly dependent on the RbBP5AS+ABM/ASH2LSPRY dimer, but not WDR5. In contrast, the catalytic activity of MLL1 SET domain was only weakly stimulated by the RbBP5_AS+ABM_/ASH2L_SPRY_ dimer and instead, its optimal activity was more dependent on WDR5. Our solution model suggests an explanation for these observations.

Highly populated models of the trimeric complex in the optimal ensemble feature a direct interaction between the WD40 domain of RbBP5 and a short peptide sequence of MLL1 located between the WIN motif and the SET domain (Fig 4E). We refer to this RbBP5 binding sequence as the RBS region of MLL1 (Fig 1A). The RBS binding surface of RbBP5-NTD consists of a number of hydrophobic residues (V249, I283, L286, V287, and I289), and residues Q273 and P253 (Fig S5D). The RBS also has an intramolecular association with the SET domain of MLL1 thereby bridging the RbBP5-NTD and the MLL1-SET domain within the complex. The MLL1-SET domain residues comprising the RBS binding interface (K3825, K3828, N3861, R3871, M3897, H3898, G3899, R3903, and F3904) are shown in Fig 5A.

**Figure 5.**
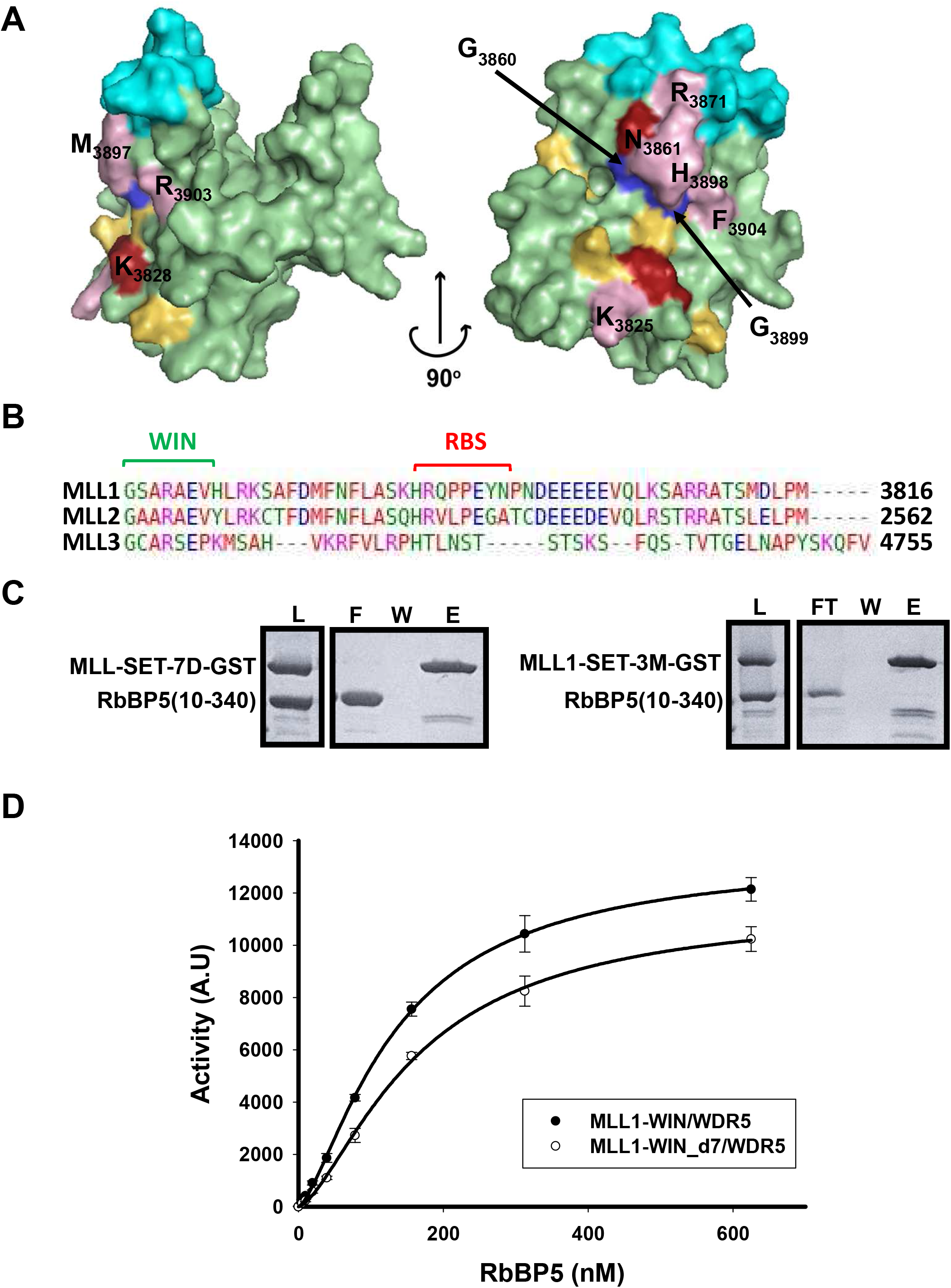
MLL1-SET domain binding interface between the RBS and RbBP5-NTD. (A) Surface representation of MLL1 SET domain (residues 3815-3969). The helix of the SET-I sub-domain is shown in cyan. Residues involved in the interface with RbBP5-NTD and the RBS are shown in pink and red. Conserved residues that are involved in the interface with the AS+AMB and ASH2L_SPRY_/RbBP5_AS+ABM_ in the MLL3 complex (32) are shown in yellow/red. Two GLY residues, that serve as the hinge points of SET-I rotation (32), are shown in blue. (B) Sequence alignment for MLL1, MLL2 and MLL3 linker residues between the WIN motif and the SET domain showing the residues that are not conserved. The WIN motif is highlighted in a green bracket. The residues within the MLL1 RBS segment are shown in a red bracket. (C) No interaction was detected between MLL1-SET-7D-GST (aa 3786-3793 deletion) and MLL1-SET-Q3787V/P3788L/Y3791G-GST (MLL1 to MLL2 mutation) mutants and RbBP5-NTD by GST pulldown experiments. Lanes are labeled as in Figure 3. (D) Deletion of the RBS region of MLL1 (MLL1-SET-7D) attenuates the catalytic activity of the trimeric complex. See also Fig S5.

Several distinct features of this putative RbBP5-MLL1 association are notable. First, in the trimeric complex the MLL1-SET domain interacts with RBS and RbBP5-NTD via a surface that significantly overlaps with the previously identified ASH2L/RbBP5 interaction surface of the SET domains of the other MLL family members (32). Further mutagenesis and computational studies (32) had suggested that this interaction surface mediates a stabilization of the flexible SET domains of MLL2,3,4 upon interaction with the ASH2L/RbBP5 dimer, which in turn, stimulated catalytic activity of the respective SET domains. We define this surface of MLL-SET domains (including MLL1) as the activation surface. A second notable feature of our MLL1 solution model is the interaction of residues N3861 and Q3867 of the SET domain activation surface with RBBP5-NTD. These two residues are unique to MLL1 and were previously shown to be incompatible with functional interactions with the ASH2L/RbBP5 dimer (32). This suggests that the MLL1 activation surface may rely on interactions with RBS and RbBP5-NTD as an alternative activation mechanism. Thirdly, the RBS segment is a unique feature of MLL1 and is not conserved in the other MLL proteins that rely more heavily on the ASH2L/RbBP5 dimer for activation (Fig 5B). According to our solution structural ensemble the RBS plays a key role in association of MLL1 with RbBP5-NTD. This is supported by both GST-pull down and gel filtration experiments. Indeed, either deletion or mutation of RBS residues of MLL1 to those of MLL2 within the MLL1-SET construct showed no interaction with RbBP5-NTD by pull down and gel filtration experiments (Fig 5C). Finally, deletion of the RBS also decreased the ability of RbBP5 to stimulate the catalytic activity of the WDR5/MLL1-WIN complex (Fig 5D).

Taken together, our structural model of the RbBP5/WDR5/MLL1-WIN trimeric complex suggests that the activation mechanism of the complex is mediated in part through the unique but likely weak interaction of RbBP5 with MLL1, which in turn stabilizes the SET-I motif of the catalytic SET domain. WDR5 serves as a hub that brings together the MLL1-SET and the WD40 repeat domain of RbBP5, thereby increasing their effective local concentrations and facilitating what is otherwise a weak (K_D_ > 1μM), but specific interaction within the trimeric complex. Hence, we hypothesized that a triumvirate of weak, but specific intermolecular interactions are required to maintain the integrity of the MLL1 minimal complex and that disruption of an individual interaction site may be sufficient to disrupt the entire functional complex. To test this hypothesis, we measured the ability of OICR-9429, a small molecule antagonist of the WDR5-WIN interaction, to disrupt the association of the RbBP5/WDR5/MLL1-WIN complex using gel filtration (Fig 6A). Pharmacological disruption of the WDR5-MLL interaction compromised the assembly of the trimeric complex (Fig 6A,B). OICR-9429 also inhibited the catalytic activity of the recombinant trimeric complex (Fig 6C). These results are consistent with our previous work showing that OICR-9429 can disrupt assembly and function of the endogenous MLL1 complex in cells (33), and similar results have been reported for MM-401, a peptide-based antagonist of MLL-WIN interaction (34, 50).

**Figure 6.**
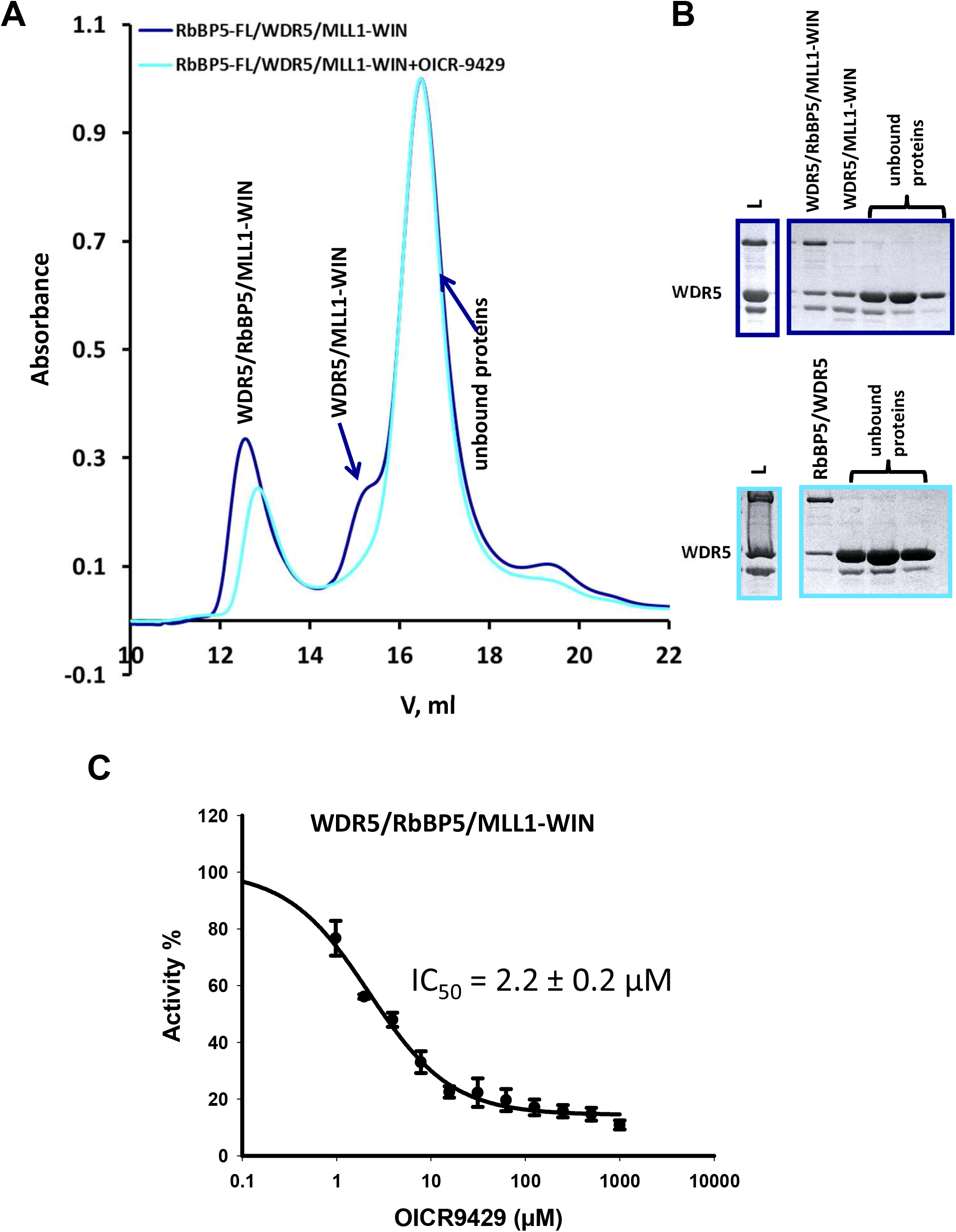
OICR-9429 attenuates the assembly of a functional trimeric complex. (A) Size exclusion chromatography of RbBP5/WDR5/MLL1-WIN (at concentrations of ~ 13 μM/12 μM/6 μM respectively) in the absence (navy) or presence of (cyan) of ~ 5-fold molar excess of OICR-9429. This compound binds to WDR5 (KD = 93 nM) (33) and competes with the MLL1 WIN motif for binding to WDR5. (B) SDS-PAGE of elution fractions (L=loaded protein mixture). Fractions containing the trimeric and WDR5/MLL1-WIN (shoulder at 15.2 ml) complexes are not recovered from the column when run in the presence of OICR-9429. (C) OICR-9429 inhibits the catalytic activity of the trimeric complex.

Our results provide the first atomic level description of a functional MLL1 complex. Our model reveals that the minimal active MLL1 complex is comprised of a series of three weak (μM) protein-protein interactions, each of which can be a site of vulnerability for disruption of the functional complex. This has important implications for development of pharmacological antagonists of the MLL1 complex and further strengthens this approach to target other multiprotein complexes that are dependent on weak, but druggable interaction sites.

## ACKNOWLEDGEMENTS

For the SAXS experiments, we gratefully acknowledge use of the SAXS Core facility of the Center for Cancer Research, National Cancer Institute (NCI). The SAXS data were collected at beamline 12-ID-B. The shared scattering beamline 12-ID-B resource is allocated under the PUP-24152 agreement between the National Cancer Institute and Argonne National Laboratory (ANL). We thank Dr. Xiaobing Zuo (ANL) for his expert support. Use of the Advanced Photon Source, a U.S. Department of Energy (DOE) Office of Science User Facility, was operated for the DOE Office of Science by Argonne National Laboratory under Contract No. DE-AC02-06CH11357. The SGC is a registered charity (number 1097737) that receives funds from AbbVie, Bayer Pharma AG, Boehringer Ingelheim, Canada Foundation for Innovation, Eshelman Institute for Innovation, Genome Canada through Ontario Genomics Institute, Innovative Medicines Initiative (EU/EFPIA) [ULTRA-DD grant no. 115766], Janssen, Merck & Co., Novartis Pharma AG, Ontario Ministry of Economic Development and Innovation, Pfizer, São Paulo Research Foundation-FAPESP, Takeda, and the Wellcome Trust.

---

**Figure S1.**
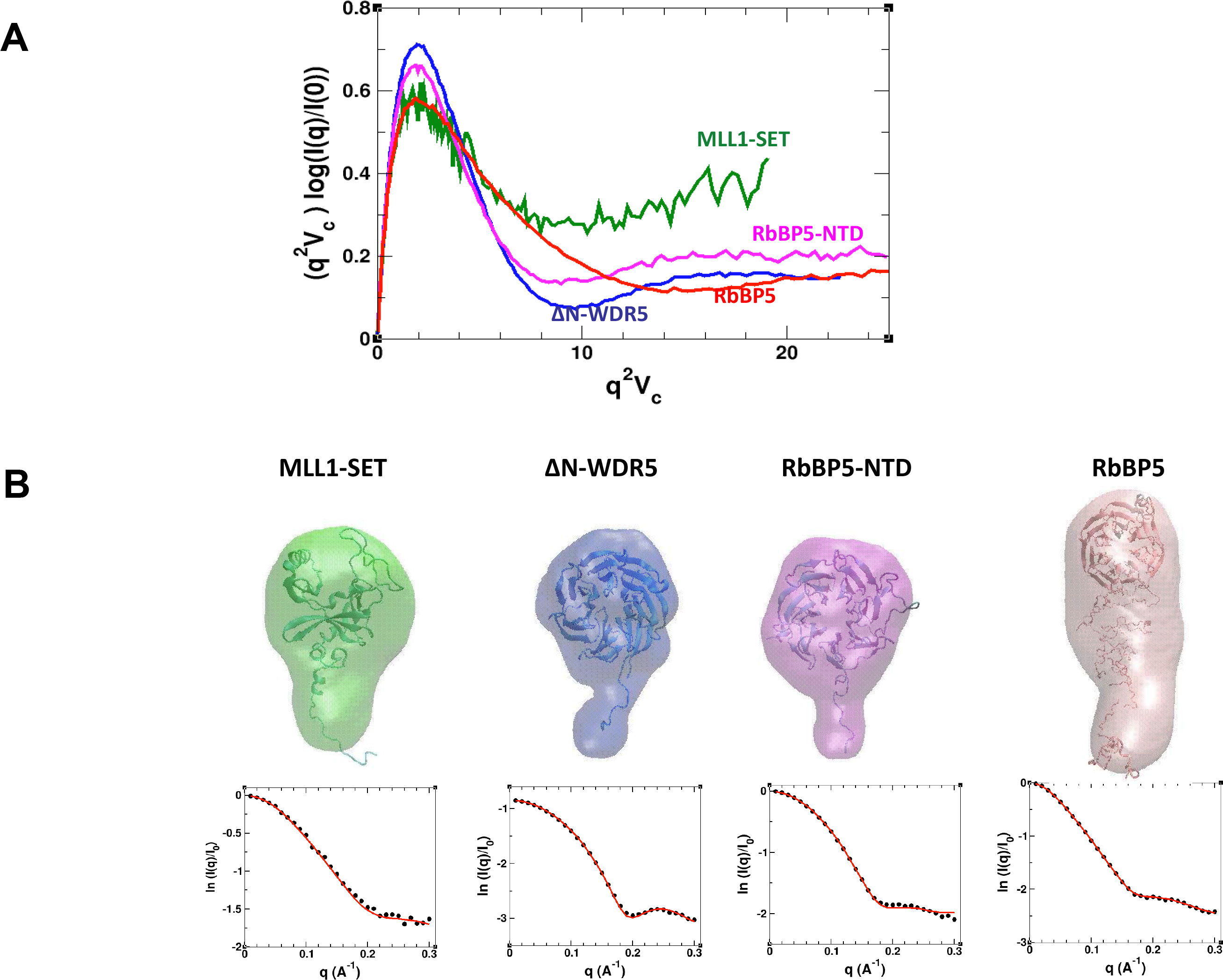
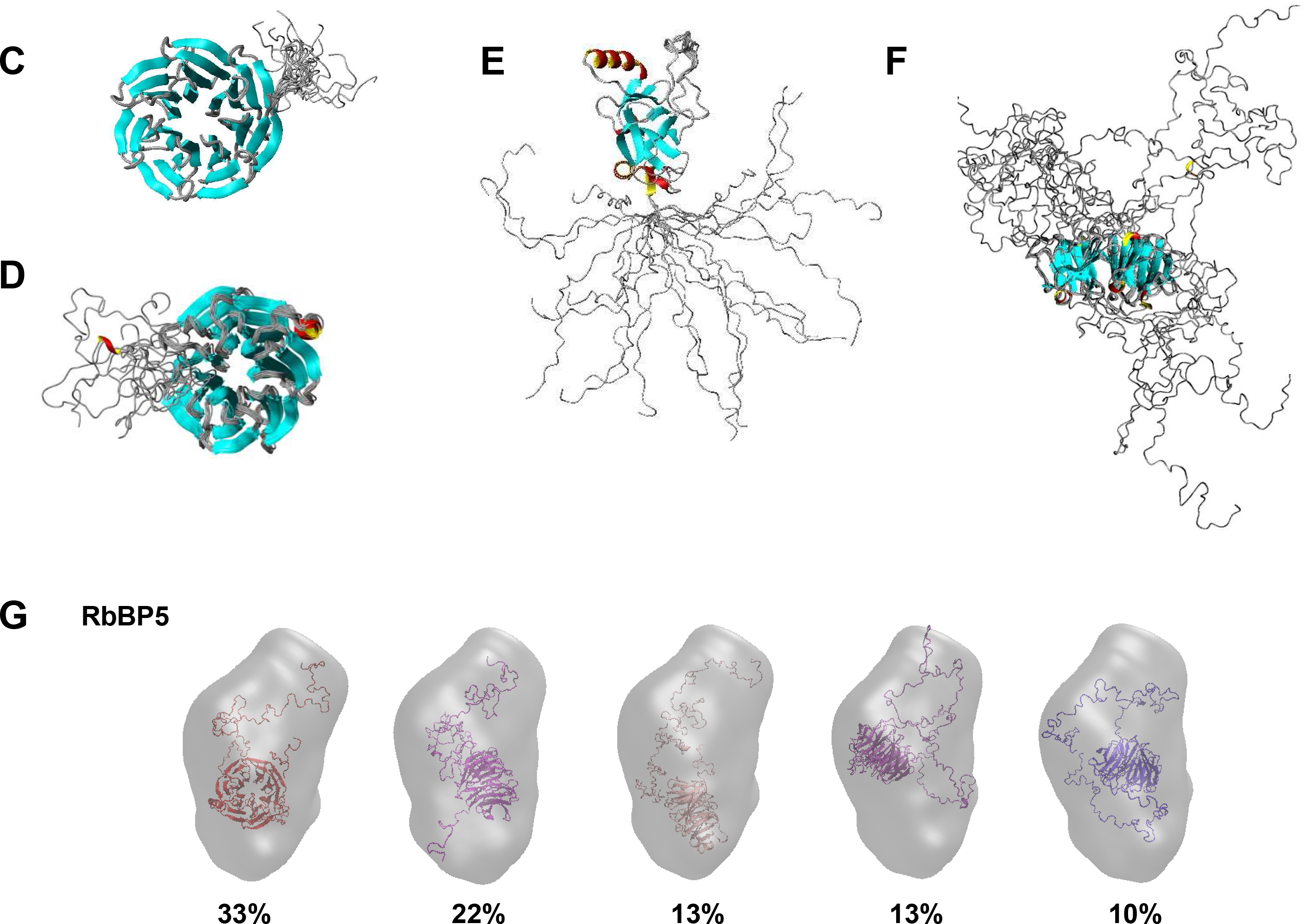
Individual components of Wdr5-RbBP5-MLL1 core complex exhibit different degrees of flexibility. (**A**) Vc-based Kratky plots of experimental SAXS data for MLL1-SET (green), ΔN-WDR5 (blue), RbBP5-NTD (magenta), and full-length RbBP5 (red). (**B**) *Top row: Ab-initio* SAXS-predicted molecular envelopes with fitted molecular models. Filtered envelopes were calculated from fifteen runs of DAMMIF. The fitted models for MLL1-SET and ΔN-WDR5 are the crystal structures (PDBID: 2W5Y and 2H9M, respectively). For RbBP5-NTD, a homology model obtained with ROSETTA is shown. For full-length RbBP5, the five most populated models of the dynamic ensemble calculated using the SES method are shown (see also panel **G**). *Lower row*: Experimental SAXS profiles (black circles) superimposed with theoretical profiles (red lines) calculated for dynamical models of MLL1-SET, ΔN-WDR5, RbBP5-NTD, and full-length RbBP5 (for details see the Materials and Methods section). Ribbon representation for the optimal ensembles of ΔN-WDR5 (**C**), RbBP5-NTD (**D**), MLL1-SET with flexible N-terminal tail (**E**), and RbBP5 (**F**). For all ensembles shown, the structured parts from different members are superimposed. (**G**) Ribbon diagrams of the five most populated species in the optimal ensemble models of RbBP5, overlaid with the average *ab-initio* SAXS-predicted molecular envelope (gray mesh).

**Figure S2.**
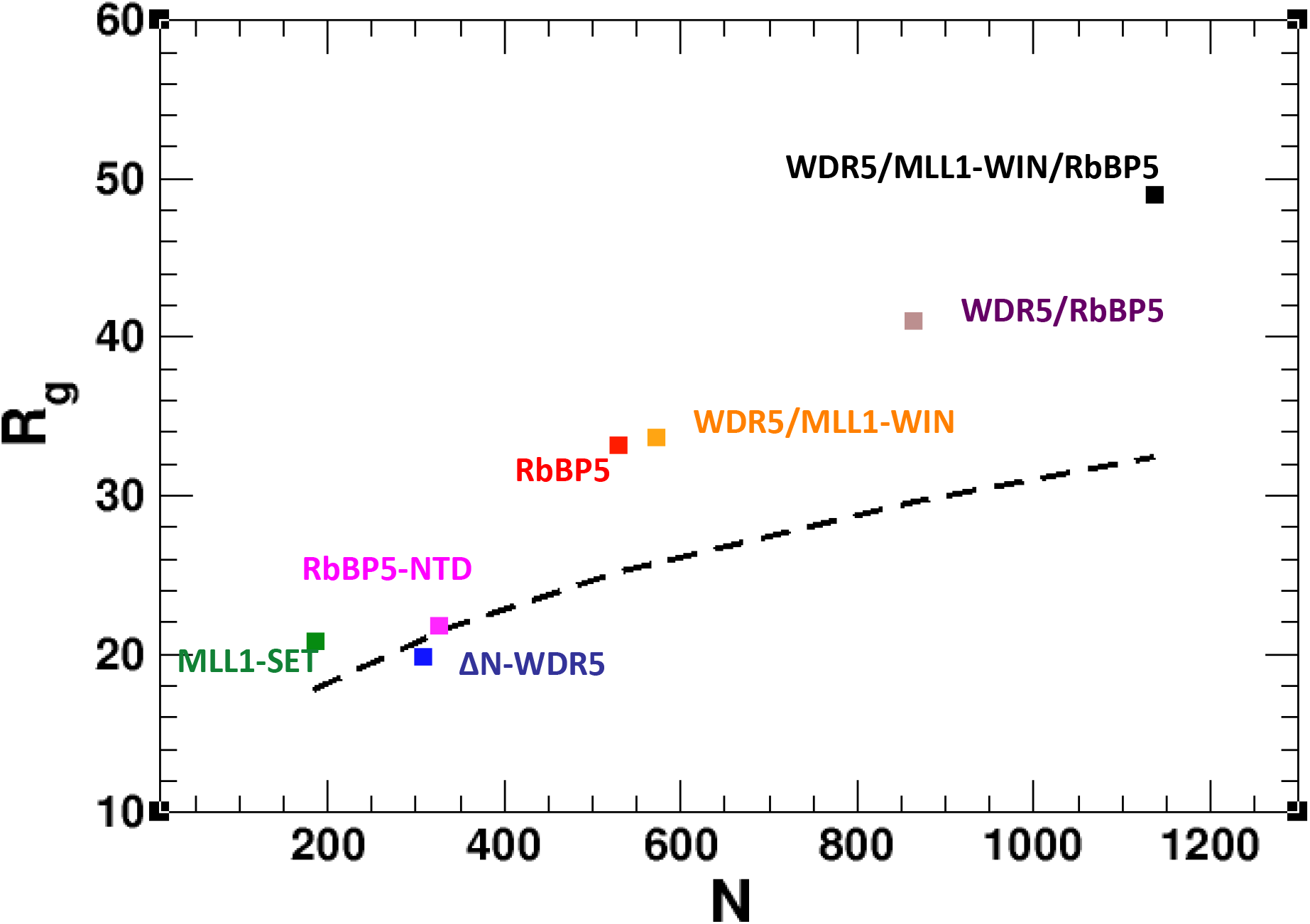
Comparison of the experimental Rg with the expected Rg for globular proteins. Experimental Rg derived from SAXS data *vs*. the number of residues (N) for MLL1-SET (green), ΔN-WDR5 (blue), RbBP5-NTD (magenta), full-length RbBP5 (red), WDR5/MLL1-WIN complex (orange), WDR5/RbBP5 complex (maroon), and WDR5/MLL1-WIN/RbBP5 complex (black). Theoretical Rg expected for a globular protein with the same molecular mass is shown by the dashed black line.

**Figure S3.**
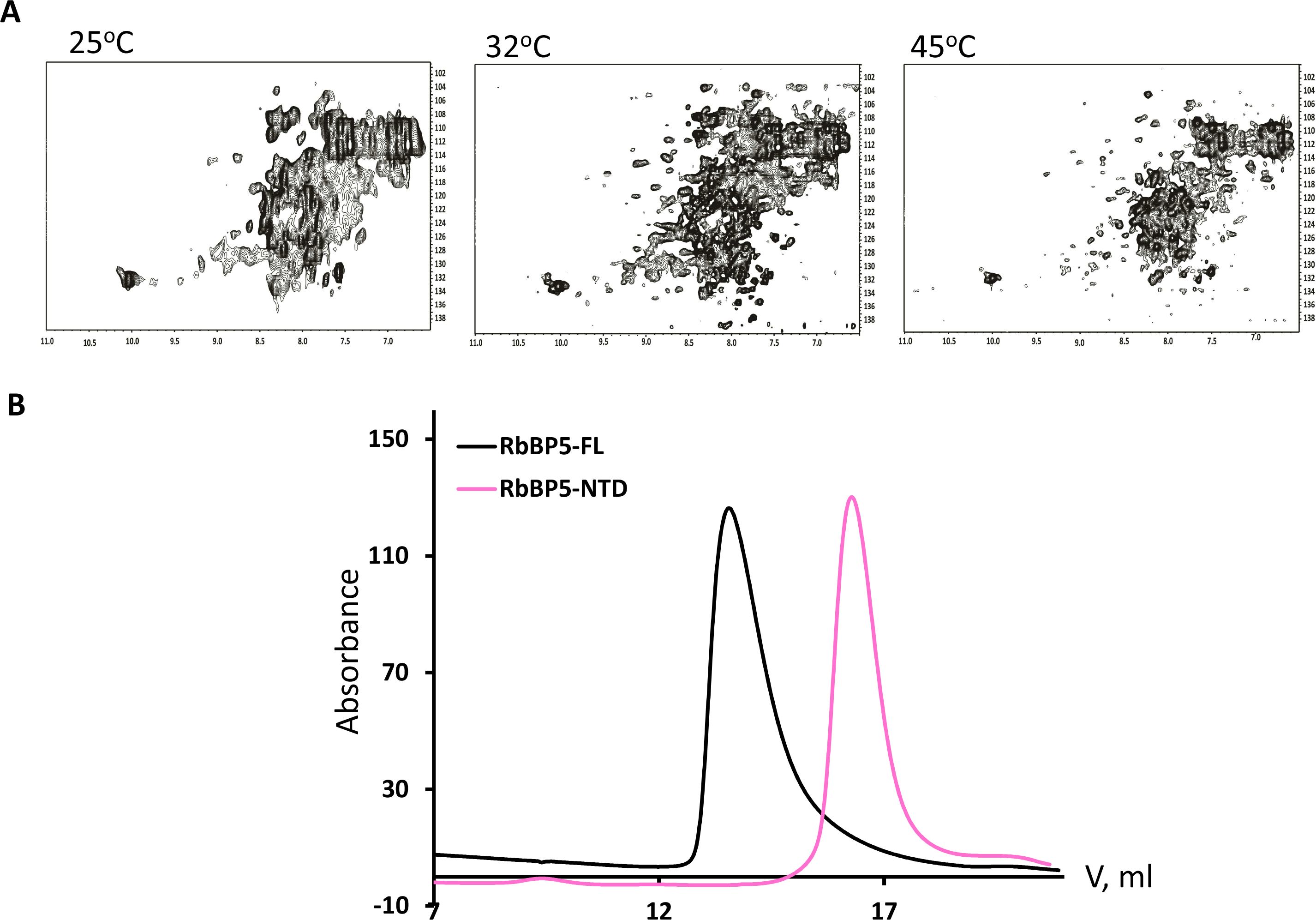
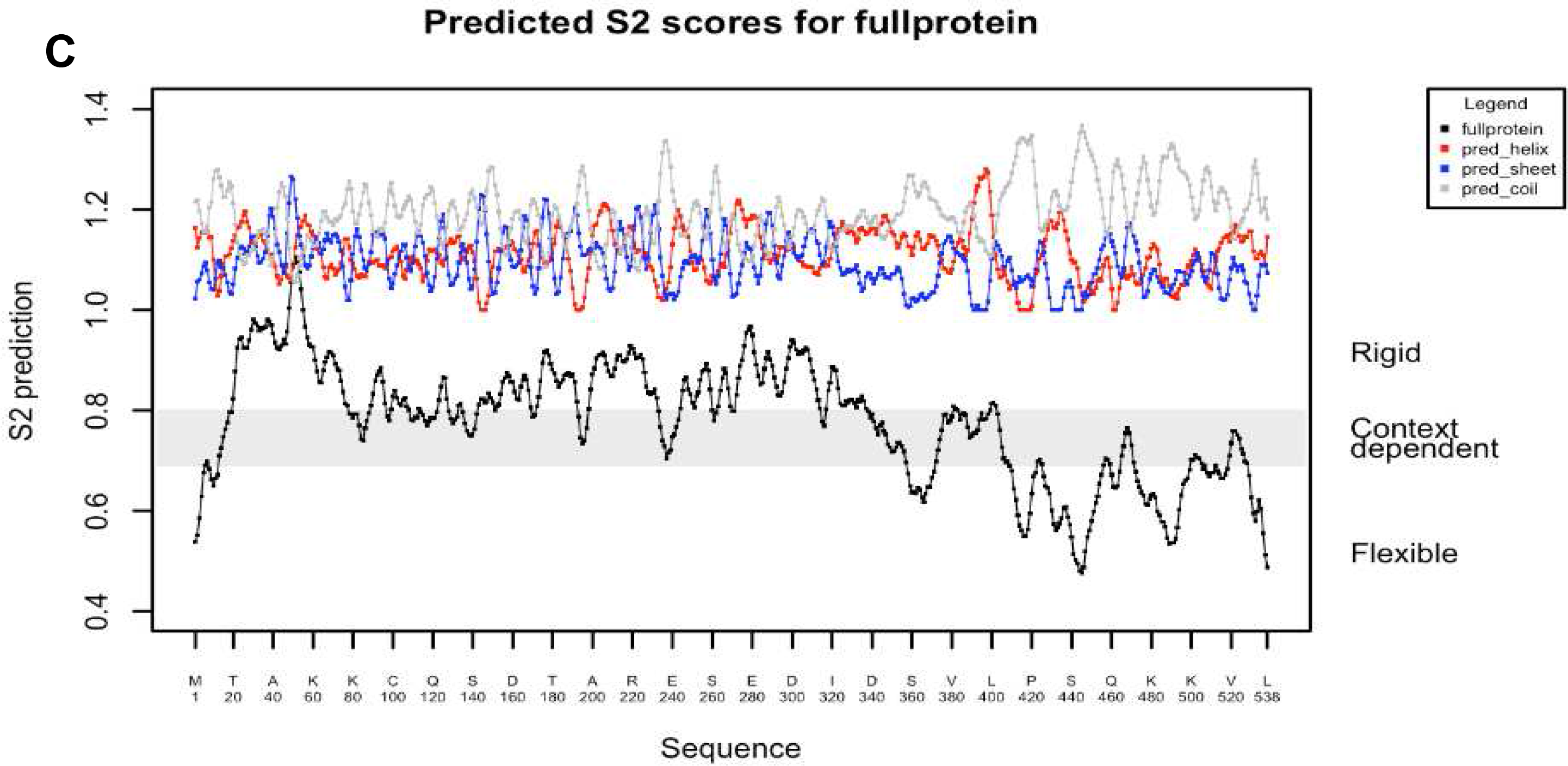
Structural characterization of full-length RbBP5. (**A**) [^1^H-^15^N]-TROSY spectra of RbBP5 collected at 25°C, 32°C and 45°C. (**B**) Gel filtration profiles of RbBP5 full-length **(black)** and RbBP5-NTD **(pink)**. RbBP5-FL runs with a higher molecular mass (MW=112.2 kDa, Ve=13.3ml) than its calculated value (MW=59.2kDa) indicating a high degree of disorder. RbBP5-NTD runs as a globular, folded domain with an elution volume (Ve=16.2 ml) corresponding to its expected molecular mass (MW=36.8kDa). (**C**) S2 order parameter for RbBP5 predicted from the protein sequence using DYNAMINE.

**Figure S4.**
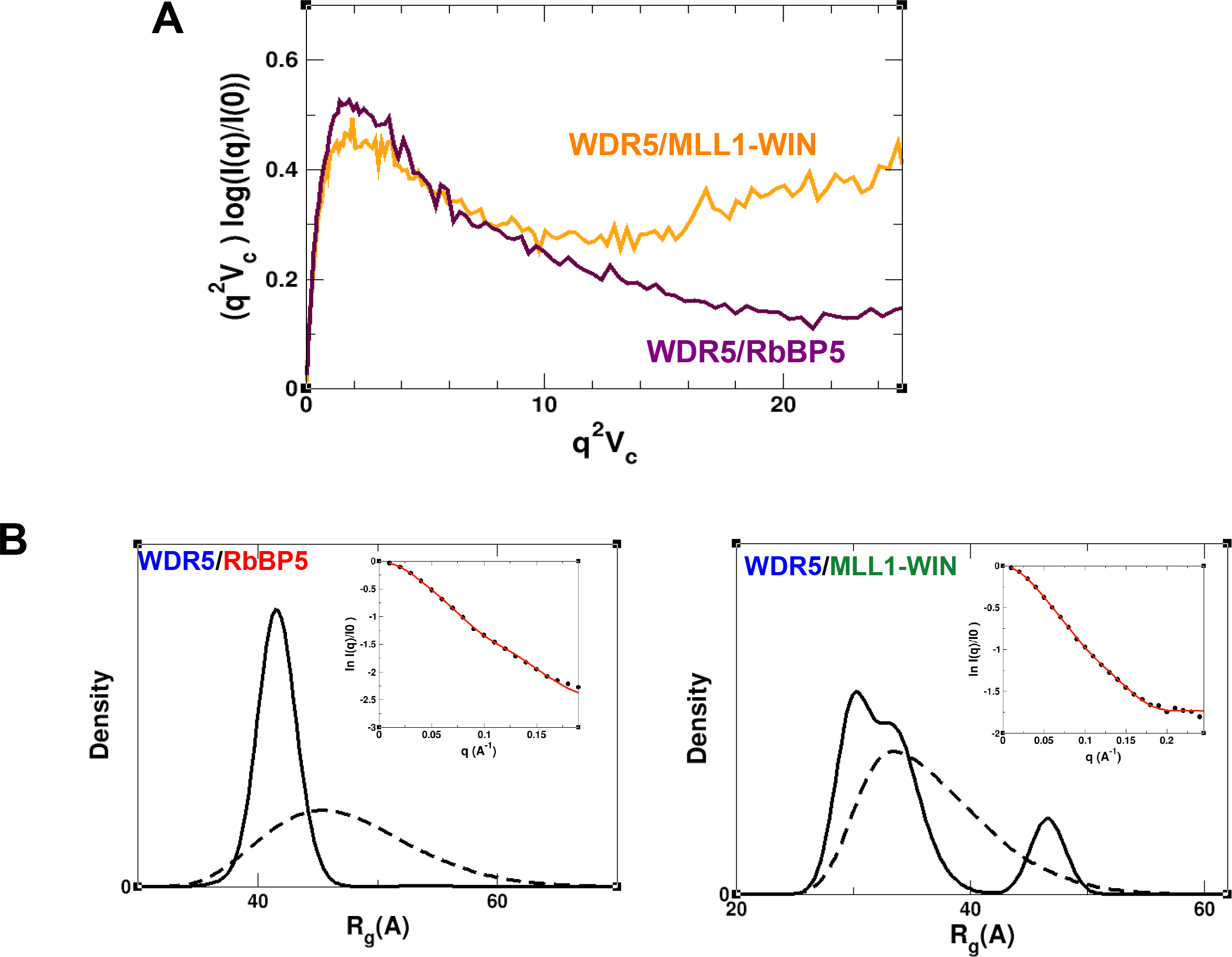
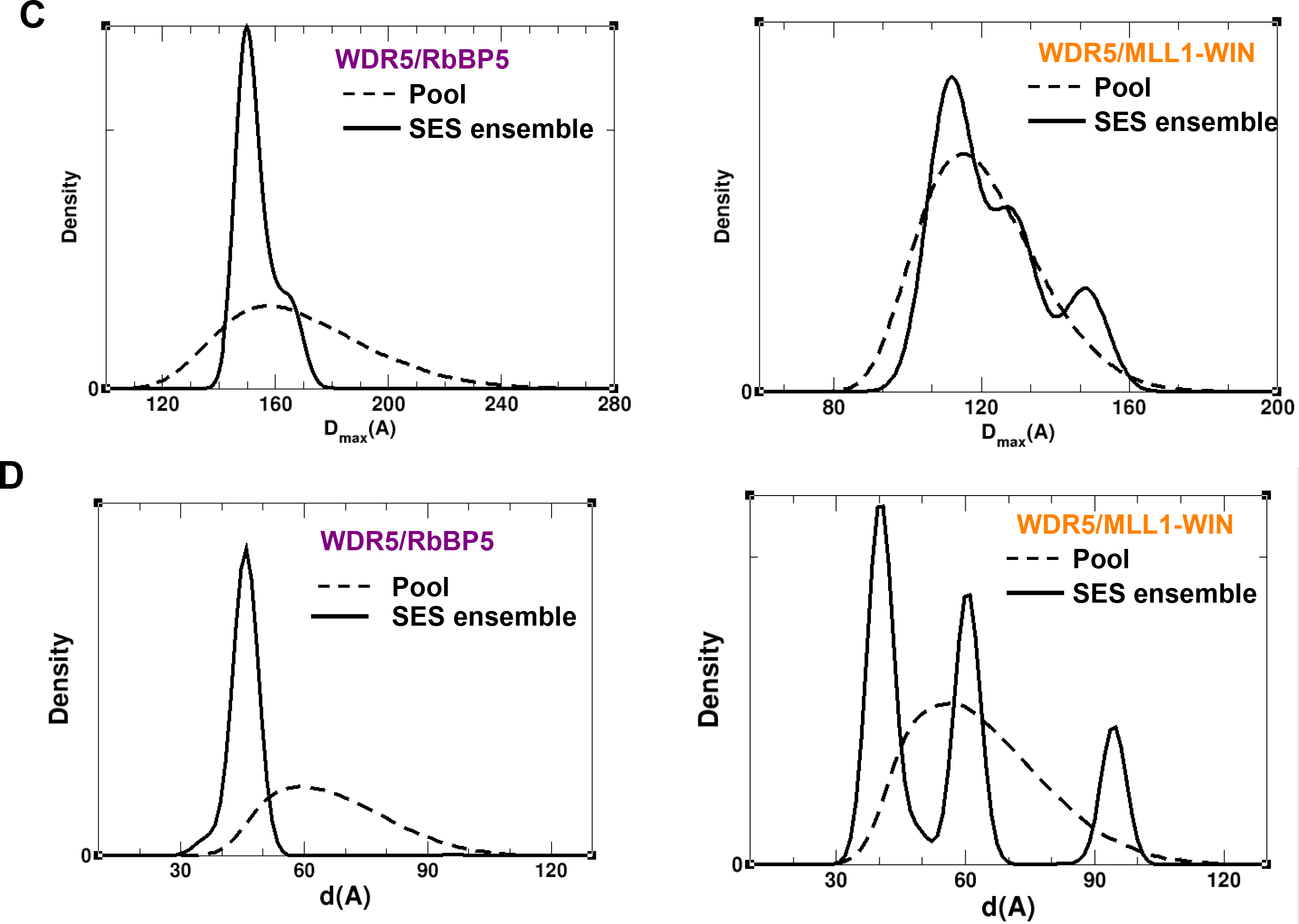
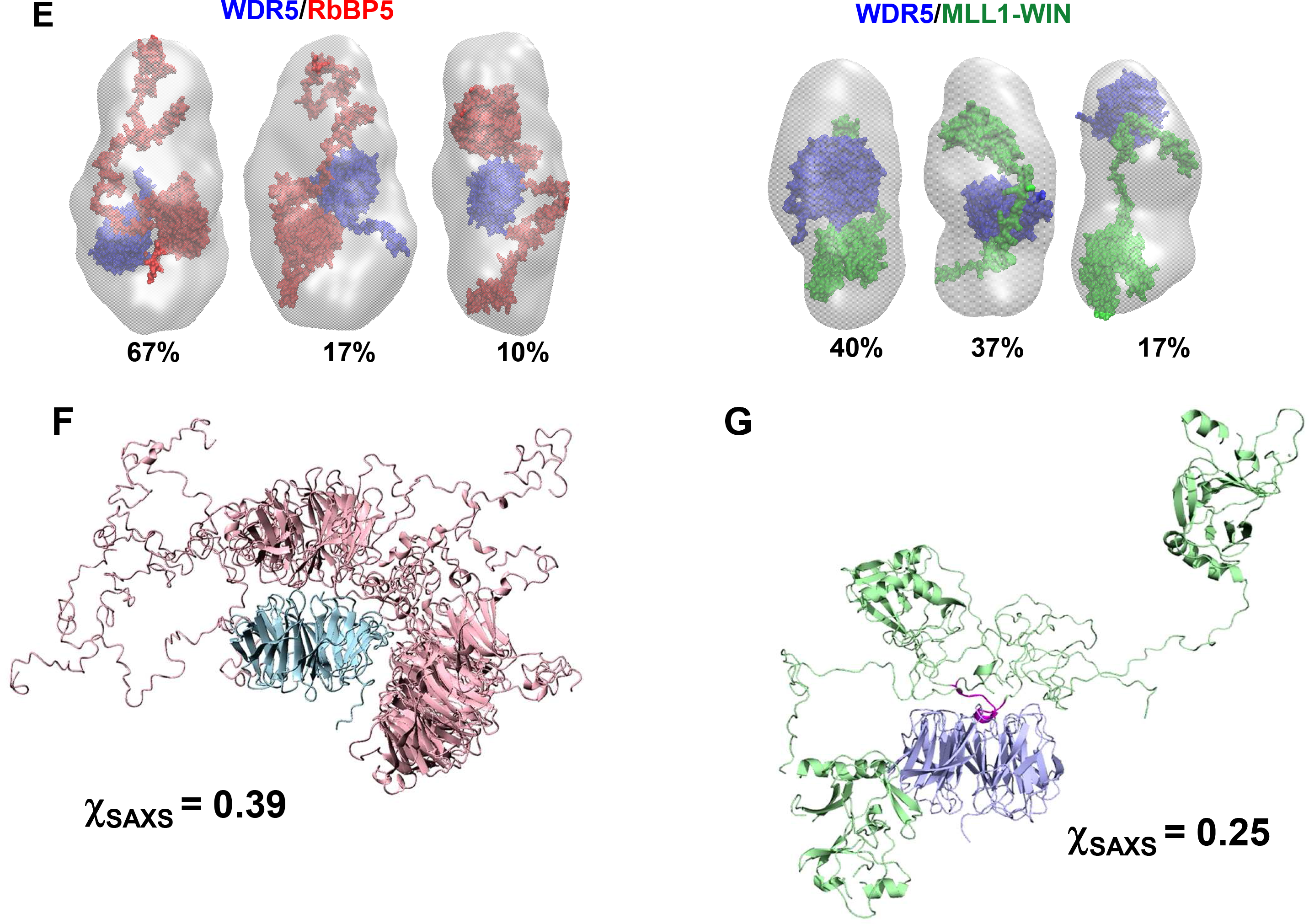
Structural characterization of WDR5/MLL1-WIN and WDR5/RbBP5 binary complexes. (**A**) Vc-based Kratky plots derived from experimental SAXS data for WDR5/MLL1-WIN (orange) and WDR5/RbBP5 complexes (maroon). (**B**) Rg distributions for the initial pool of random structures (dashed), and for the selected SES ensemble (solid). Experimental SAXS profiles (black circles) in a match with theoretical profiles (red lines) averaged over the SES ensemble are shown (inset). (**C**) D_max_ distribution for WDR5/MLL1-WIN (left) and WDR5/RbBP5 (right). The distribution for the initial pool of random structures (dashed lines) and for the selected SES ensemble (solid lines) are shown. WDR5/RbBP5 favors structures that are more condensed than fully extended, while WDR5/MLL1-WIN has a higher degree of flexibility and exhibits a population of fully extended structures. (**D**) Distribution of the distance between centers of mass of WDR5 and globular RbBP5 (right panel), and WDR5 and MLL1-SET (left panel) domains in the optimal SES ensembles of the binary complexes. (**E**) Surface representation of models for the binary complex overlaid with the *ab-initio* SAXS-predicted molecular envelopes (gray mesh) for WDR5/RbBP5 (left) and WDR5/MLL1-WIN (right). The three most populated models of the optimal SES ensemble are shown. WDR5, MLL1, and RbBP5 are colored in blue, green and red, respectively (**F-G**) Ribbon diagram of the representative models of the optimal ensemble of WDR5/RbBP5 (**F**) and WDR5/MLL1-WIN (**G**). WDR5, RbBP5 and MLL1 are colored in blue, pink, and green, respectively. The WIN motif of MLL1 is shown in magenta. WDR5 structured domain of different members of an ensemble are superimposed. The goodness-of-fit of the optimal ensemble to SAXS data is shown by the χ_SAXS_ value.

**Figure S5.**
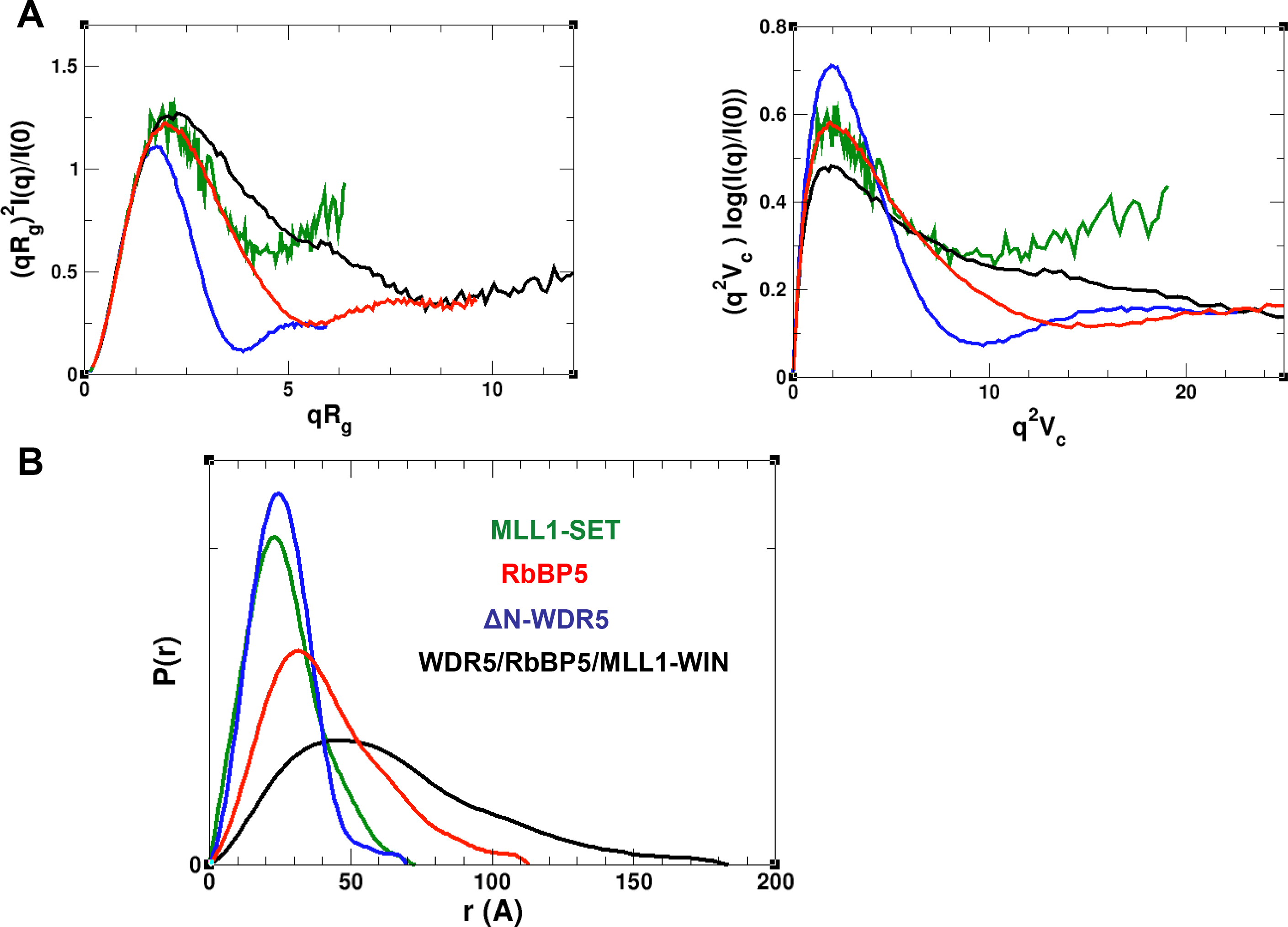
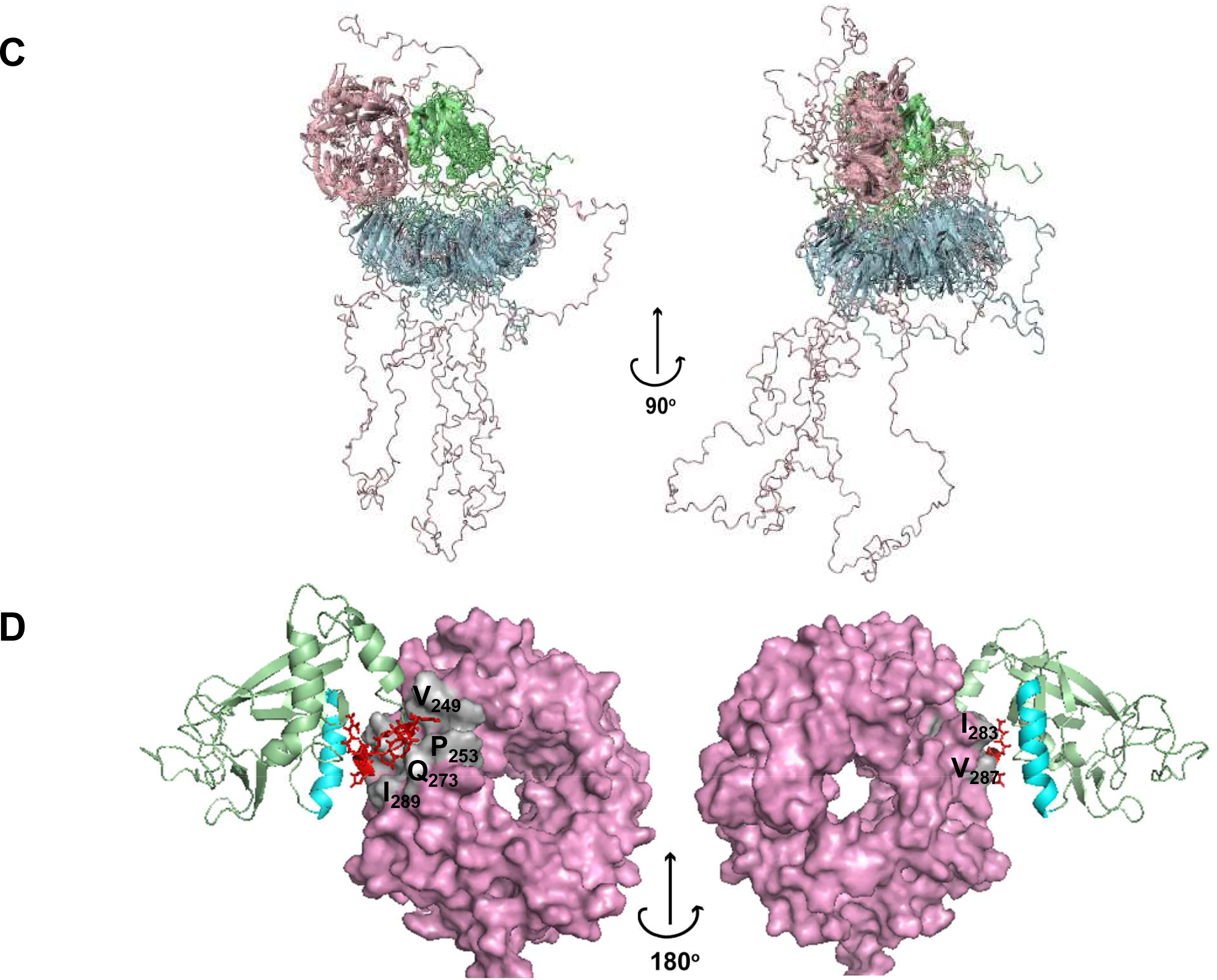
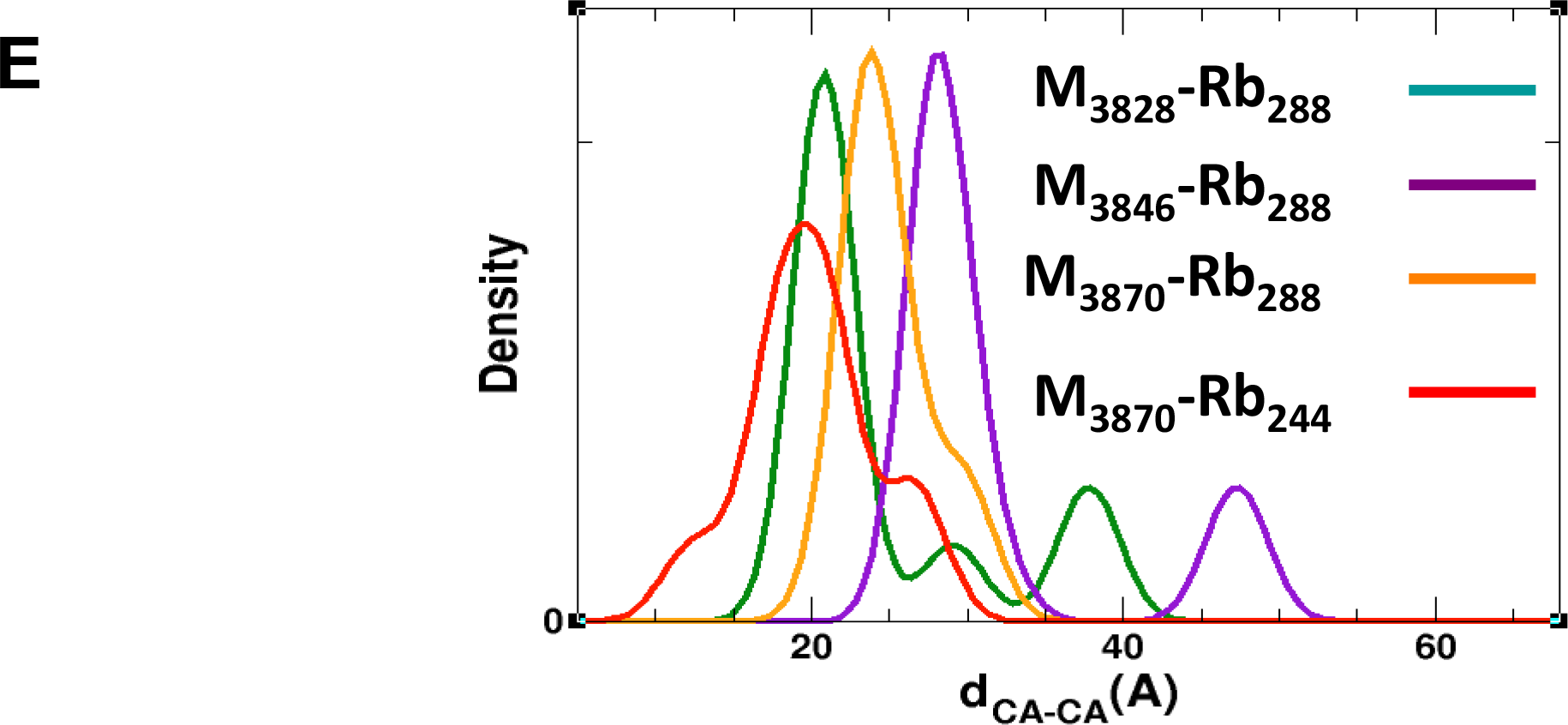
Structural characterization of trimeric WDR5/RbBP5/MLL1-WIN complex. (**A**) Rg-based (left panel) and Vc-based (right panel) Kratky plots of experimental SAXS data for the WDR5/RbBP5/MLL1-WIN complex (black), and its components, ΔN-WDR5 (blue), MLL1-SET (green), and full-length RbBP5 (red line). SAXS data indicate a high degree of flexibility for the complex. (**B**) Normalized pair distance distribution function P(**r**) calculated from experimental SAXS data with GNOM. (**C**) Ribbon diagram of six representative models of the optimal ensemble of the complex. These models comprise 86% of the ensemble. WDR5, RbBP5 and MLL1 are colored in blue, pink, and green, respectively. Structured N-terminal domain of RbBP5 and SET domain of MLL1 from different members of the ensemble are superimposed. (**D**) Molecular model of RbBP5-NTD/MLL1-SET complex. RbBP5-NTD surface is colored in pink. MLL1-SET is shown as a ribbon diagram in pale green. The helix of the SET-I sub-domain is highlighted in cyan. The RBS (MLL1_3785-3792_) residues are shown in red. The RbBP5 residues that are in contact with MLL1-SET are shown in gray. (**E**) Ensemble distribution of four X-links between RbBP5-NTD and MLL1-SET domains.

**Figure S6.**
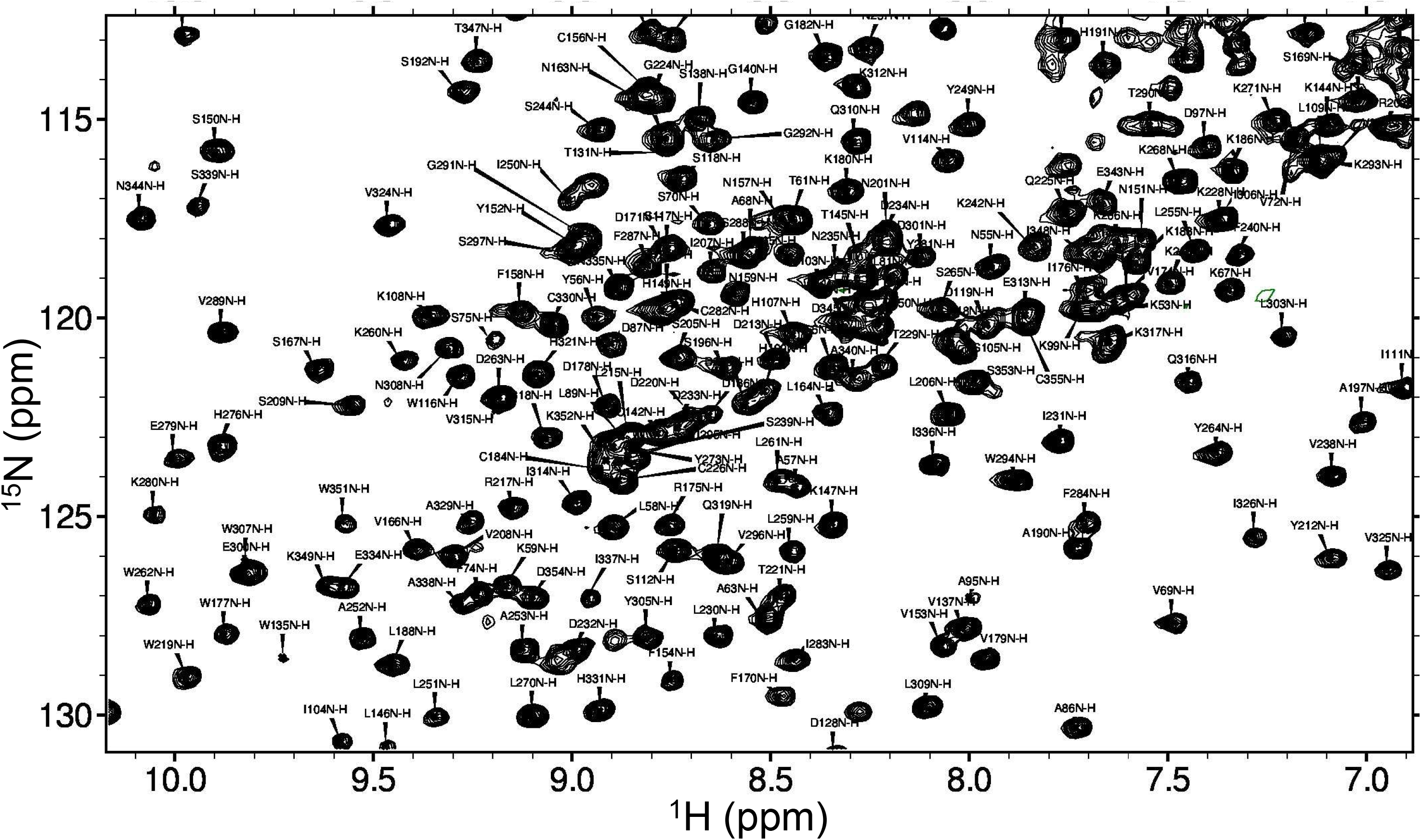
Portion of an [^1^H-^15^N]-TROSY spectrum of ΔN-WDR5. Peaks are labeled with resonance assignments – 254 backbone amides were assigned in the construct.

## Detailed Materials and Methods

### Cloning of MLL1, WDR5 and RbBP5 constructs

The coding regions for human MLL1-SET (aa3785-3969) and MLL1-WIN (aa3745-3969) were PCR-amplified and sub-cloned into the pET28GST-LIC vector (GeneBank ID: EF456739). The mutants, MLL1-SET-7D (Δ3786-3793) and MLL1-SET _Q3787V/P3788L/Y3379G, were generated from the wild-type clone using the QuikChange PCR mutagenesis kit (Agilent). RbBP5 of different lengths (aa 10-340, 320-410, 340-538, 1-538) and WDR5 (aa 24-334) were sub-cloned into pET28-MHL vector (GeneBank ID: EF456738). For expression of dimeric complexes and reconstitution of trimeric MLL1 complex, full-length WDR5 and MLL1-WIN, WDR5 and RbBP5 were cloned into pFastBac Dual expression vector (Thermo Fisher Scientific), respectively, with MLL1-WIN and RbBP5 tagged with N-terminal hexa His-tag.

### Protein expression and purification

Human MLL1 constructs containing residues 3745-3969 (MLL1-WIN) and 3785-3969 (MLL1-SET), as well as the WD-40 repeat region of WDR5 (residues 24-334), the N– and C-terminus of RbBP5 (RbBP5-NTD, RbBP5-CTD), and full-length RbBP5 were individually expressed in *E. Coli* (BL21(DE3) codon plus RIL, Agilent). The MLL1 proteins were expressed as N-terminal GST fusions and purified on a GST-bind (Novagen) column according to manufacturer’s instruction. The GST tag was cleaved off by incubating the resin-bound fusion protein with thrombin (Sigma) at 4 C overnight. The eluted MLL1 proteins were passed through a gel filtration column (Superdex 200, GE Healthcare) pre-equilibrated with 20 mM Tris-HCl (pH 7.5), 500 mM NaCl. WDR5 and RbBP5 proteins were expressed as N-terminal Hexa-His fusions and purified on Nickle-chelating column (GE Healthcare). After elution and removing His-tag by incubation with TEV protease at 4 °C overnight, the proteins were subjected to a gel filtration column Superdex 200, GE Healthcare) pre-equilibrated with 20 mM Tris-HCl (pH 8.0), 500 mM NaCl (for RbBP5) and 20 mM PIPES (pH 6.5), 250 mM NaCl (for WDR5), respectively.

The dimeric and trimeric complexes of MLL1 used for SAXS and cross-linking studies were expressed in Sf9 cells. The dimeric complex of WDR5-MLL1-WIN and WDR5-RbBP5 were purified by TALON affinity column (Clontech) followed by a gel filtration column (Superdex 200, GE Healthcare) preequilibrated with 20 mM BisTris propane (pH 7.0), 250 mM NaCl. The fractions containing the two dimeric complexes were collected separately and used for SAXS data collection. The two dimeric complexes were mixed and incubated on ice for 2 hours and further purified on a gel filtration column (Superdex 200, GE Healthcare) pre-equilibrated with 20 mM BisTris propane (pH 7.0), 250 mM NaCl. The fractions containing the trimeric complex were collected and used for SAXS data collection and cross-linking experiments.

### SAXS data collection and analysis

SAXS measurements were carried out at the beamline 12-ID-B of the Advanced Photon Source, Argonne National Laboratory. The energy of the X-ray beam was 14 Kev (wavelength λ=0. 8856 Å), and two setups (small-and wide-angle X-ray scattering, SAXS and WAXS) were used simultaneously in which the sample to Pilatus 2M detector distance were adjusted to achieve scattering *q* values of 0.006 < *q* < 2.6Å^-1^, where q = (4π/λ)sinθ, and 2θ is the scattering angle. Thirty two-dimensional images were recorded for each buffer or sample solutions using a flow cell, with the accumulated exposure time of 0.8-2 seconds to reduce radiation damage and obtain good statistics. No radiation damage was observed as confirmed by the absence of systematic signal changes in sequentially collected X-ray scattering images. The 2D images were corrected and reduced to 1D scattering profiles using the Matlab software package at the beamlines. The 1D SAXS profiles were grouped by sample and averaged. The scattering profile of the protein was calculated by subtracting the background buffer contribution from the sample-buffer profile using the program PRIMUS (ATSAS package, EMBL) (1). Concentration series measurements for each sample were carried out to remove the scattering contribution due to inter-particle interactions and to extrapolate the data to infinite dilution. The protein concentration ranges used for MLL1-WIN/WDR5, RbBP5/WDR5, and MLL1-WIN/WDR5/RbBP5 were 14-28 μM, 10-36 μM and 9.6-46.5 μM, respectively. These concentrations are >10 fold of the dissociation constants for each binary interaction of the dimers. Guinier analysis and the experimental radius of gyration *(R_g_)* estimation from the data of infinite dilution were performed using PRIMUS. The pair distance distribution function (PDDF), p(**r**), and the maximum dimension of the protein, *D*_max_, in real space was calculated with the indirect Fourier transform using program GNOM (2). To avoid underestimation of the molecular dimension and consequent distortion in low resolution structural reconstruction, the parameter *D*_max_, the upper end of distance *r*, was chosen such that the resulting PDDF has a short, near zero-values tail at large *r*. The Rg from P(**r**) analysis was also reported. The Volume of correlation (3), V_c_, was calculated using in-house script. The molecular weights were estimated using V_c_ (3) in q range of 0 < *q* < 0.3 Å^-1^. Fifteen *ab-initio* shape reconstructions (molecular envelopes) were generated using DAMMIF (4) and averaged with DAMAVER (5). The structural models were superimposed and overlaid with the averaged envelop using SUPCOMP (6). The theoretical scattering intensity of a structural model was calculated and fitted to the experimental scattering intensity using CRYSOL (7) and FoXS (8)programs.

### Chemical cross-linking mass spectrometry

The reconstituted trimer complex of WDR5, RbBP5 and MLL1-SET was cross-linked at a concentration between 12 and 16 μM with 1 mM of isotopically coded disuccinimidyl suberate (DSS-d_0_,DSS-d_12_) for 30 minutes at 37 °C while shaking at 500 rpm on a Thermomixer (Eppendorf) as previously described (9). Samples were quenched with 50 mM NH_4_HCO_3_ for 20 minutes at 37 °C and evaporated to dryness in a vacuum centrifuge. The dried pellets were then dissolved in 50 μL of 8 M urea, reduced with 2.5 mM Tris(2-carboxyethyl)phosphine (Pierce) for 30 minutes at 37 °C and alkylated with 5 mM iodoacetamide (Sigma-Aldrich) for 30 minutes at room temperature, in the dark. Digestion was carried out after diluting urea to 5 M with 50 mM NH_4_HCO_3_ and adding 1% (w/w) LysC protease (Wako Chemicals) for 2 hours at 37 °C and subsequently diluting to 1 M urea with 50 mM NH_4_HCO_3_ and further adding 2% (w/w) trypsin (Promega) for 14 hours at 37 °C. Protein digestion was stopped by acidification with 1% (v/v) formic acid. Digested peptides were purified using Sep-Pak C18 cartridges (Waters) according to the manufacturer’s protocol. Cross-linked peptides were enriched by peptide size-exclusion chromatography (SEC) as previously described (9). SEC fractions were then reconstituted in 5% acetonitrile and 0.1% formic acid and analysed in duplicates on a LC (Easy-nLC 300) coupled to a mass spectrometer (Orbitrap LTQ XL). Analytes were separated on self-packed New Objective PicoFrit columns (11 cm x 0.075 mm I.D.) containing Magic C18 material (Michrom, 3 um particle size, 200 Å pore size) over a 60-min gradient from 7% to 35% acetonitrile at a flow rate of 300 nL/min. The mass spectrometer was operated in data-dependent acquisition (DDA) mode with MS acquisition in the Orbitrap analyzer at 60,000 resolution and MS/MS acquisition in the linear ion trap at normal resolution after collision-induced dissociation. DDA was set up to select up to five most abundant precursors with a charge state of +3 or higher (9). MS data were converted to mzXML format with msConvert (10) and searched with xQuest/xProphet (11) against a database containing the FASTA sequences of the analysed proteins and relative decoy sequences. Cross-linked peptides were identified with a minimal length of 5 amino acids and at least four bond cleavages or three adjacent ones per peptide. Validated cross-linked peptides had a total ion current of total ion current explained higher that 0.1 and xQuest score higher than 20. Figures were prepared with xiNET (12).

### Structural characterization using SAXS data

The SAXS data indicate that the trimeric complex and its sub-complexes, as well as individual molecules MLL1-SET and RbBP5, are flexible molecular systems in solution. Thus we take an ensemble approach for structural characterization of these systems by utilizing SES protocol (13). The strategy on which SES method is based consists of two main steps: 1) generate the initial ensemble of conformations in order to approximate the conformational space available for a system in solution; 2) find optimal weight *w_k_* for each conformation *k* from the initial ensemble that minimizes discrepancy

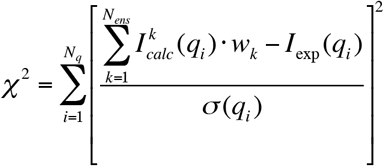

were *I_exp_(**q**)* is the experimental scattering intensity, *N_q_* is number of experimental points, *σ(**q**)* is the experimental error, and 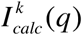 is scattering intensity predicted for *k*th conformation, and of *N_ens_* is number of conformations in the initial ensemble.

Multi-orthogonal matching pursuit (13) is used to find possible ensembles on step 2, and optimal ensemble size was select using *l*-curve. The optimal weights *w_k_* were then obtained by averaging over top solutions with similar*χ*^2^.

### Generation of the structural ensembles

#### MLL1-SET

The high degree of flexibility observed for MLL1-SET sample originate from inherent flexibility of SET domain and 28 aa long disordered N-terminal tail. We used all-atom molecular dynamics simulations to generate initial ensemble of conformers. We used all atom MD simulations to generate a trajectory started from the known crystal structure of the MLL1 SET domain with the cofactor product AdoHcy (14) (PDB id: 2W5Y). After minimization and equilibration a productive run was continued for 70 ns. Theoretical scattering profiles in the *q* range 0 < *q* < 0.3 Å^-1^ for 7,000 frames taken from the trajectory were calculated using CRYSOL.

#### RbBP5-NTD

The homology model of the N-terminal domain of RBBP5, RbBP5_24-340_, generated by ROBETTA (15), was used as starting structure for all atom MD simulations of RbBP5-NTD domain. MD trajectory of 20ns was generated and theoretical scattering profiles in the *q* range 0 < *q* < 0.3 Å^-1^ for 2000 frames taken from the trajectory were calculated using FoXs (8). The domain keeps its structure along the trajectory within 3.5 Å of backbone r.m.s.d. to the initial homology model. The calculated scattering curves were averaged over the entire ensemble of structures using the optimal weights for each ensemble member obtained with SES method, and this average profile was compared with the experimental scattering data.

#### RbBP5

The initial ensemble for SES analysis of full length RbBP5 was generated using RANCH (16) program using homology model for the WD40 domain and assuming the N-terminal RbBP5_1-24_ and C-terminal RbBP5_325-538_ regions to be disordered. The ensemble consists of 20,000 models with random conformation of the flexible regions. Theoretical scattering profile for each member of the ensemble was calculated in the *q* range 0 < *q* < 0.3 Å^-1^ using CRYSOL.

#### WDR5/MLL1-WIN

A pool of possible conformations of the dimeric complex was generated assuming the interaction between WDR5 and WIN peptide of MLL1 and utilizing known crystal structure of the WDR5/WIN complex (17)(PDB id: 3EMH). 30,000 random configurations of rigid SET domain tethered to WDR5 by WIN motif and flexible linker consisting or 46 amino acids, MLL1_3771-3817_, was generated using RANCH. Theoretical scattering profiles were calculated in the *q* range 0 < *q* < 0.25 Å^-1^ using CRYSOL.

#### WDR5/RbBP5

We used the model of full length RbBP5 described above to generate ensemble of possible conformations of the WDR5/RbBP5 complex with RANCH. We assumed that RbBP5 interacts with WDR5 by WBM motif as in the know crystal structure of sub-complex WDR5/WBM (18)(PDB ID:2XL2), so that the WD40 domain of RbBP5 and the WDR5/WBM sub-complex, both considered to be rigid in the simulations, are connected by flexible linker consisting of 48 residues, RbBP5_325-372_.

Theoretical scattering profile for each of 30,000 generated random conformations of the WDR5/RbBP5 complex was calculated in the *q* range 0 < *q* < 0.20 Å^-1^ using CRYSOL.

#### WDR5/RbBP5/MLL1-WIN

The initial ensemble of possible conformations for the trimeric complex was obtained in three steps. (**i**) On the first step, a rigid-body modeling of the complex using CORAL (19)was performed. The known ordered regions of the complex, which are assumed to be rigid on this step of simulations, consist of the known structures of MLL1 SET domain (14), sub-complex WIN/WDR5/WBM (20)(PDB id: 3P4F), and a homology model of WD40 domain of RbBP5. The rest of the complex segments (~31% of all residues) are assumed to be flexible and are modeled by chains of dummy residues. The experimental inter-domain cross-links data were taken onto account in the CORAL calculations by introducing six C_α_ – C_α_ distance restraints (see Table S1) with upper bound of 30 Å. CORAL tries to build a single conformation of the complex that fits SAXS data under the imposed constraints. Performing multiple CORAL runs we generated a number of different conformations of the complex that fit SAXS data with *χ_SAXS_* ~ 0.9. Although the obtained conformations have different interdomain arrangements the relative position of the SET domain and WD40 domain of RbBP5 is well defined and suggests the interaction of these domains in the complex. (ii) On the second step we “refined” the best CORAL models by carrying out all-atom molecular dynamic simulations. The initial conformation for MD simulations was constructed from CORAL model by building an all-atom reconstruction model using PULCHRA (21). A 20 ns MD trajectory was generated at T = 300 K. (iii) On the third step, we used coarse-grained MD simulations to generate a pool of possible conformations of trimetric complex that are consistent with known inter-molecular binary interactions and cross-links.

The structures from the step 2 were used to derive native contact map of quasi-rigid regions of the complex, which determines the nonbonded part of the Go-like potential. The quasi-rigid regions include residues WDR5_38-330_, RbBP5_29-320_, MLL1_3816-3969_, MLL1_3761-3767_, MLL1_3785-3792_, and RbBP5_374-379_ that correspond to the WD40 domains of WDR5 and RbBP5, SET domain of MLL1, WIN motif, RBS, and WBM, respectively. We have generated eight 300ns MD trajectories at T=300K. Then 24,000 conformations were saved and used as initial ensemble for fitting to SAXS data by SES method. Theoretical scattering profiles for each conformation in the ensemble were calculated in the *q* range 0 < *q* < 0.23 Å^-1^ using FoXS.

### All-atom molecular dynamics simulations

A modified Generalized Born implicit solvent model (22) was exploited in the MD simulations in order to accelerate sampling of the conformational space for each of the systems. All simulations used an integration step of 2 fs with fixed bonds between hydrogen atoms and heavy atoms. Temperature was controlled by carrying out Langevine dynamics with damping coefficient set to 2 *ps*^-1^. The cut-off for non-bonded Lennard-Jones and electrostatic interactions were set to 18 Å. Ionic strength was set to 0.15M. All simulations were performed using NAMD 2.9 code (23) with the AMBER Parm99SB parameter set (24). For residues that coordinate Zn ions a Zinc AMBER Force Field (25) was used.

### Coarse-grained molecular dynamics simulations

We used a coarse-grained model of RbBP5/MLL1-WIN/WDR5 protein complex in order to enhance the sampling efficiency in the conformational space of the complex. In this model, amino acid residues in the proteins are represented as single beads located at their C_α_ positions and interacting via appropriate bonding, bending, torsion-angle, and non-bonding potential. A Gō-like model of Clementi and Onuchic (26)was employed to maintain the structured, globular domains as quasi-rigid in the simulation. For flexible regions, we adopt simple model in which adjacent amino acids beads are joined together into a polymer chain by means of virtual bond and angle interactions with a quadratic potential.

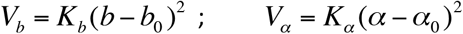

with the constants *K_b_* = 50 *kcal/mol* and *K_α_* = 1.75 *kcal/mol* and the equilibrium values *b_0_* = 3.8 Å and *α_0_* = 112° for bonds and angles, respectively. The excluded volume between nonbonded beads was treated with pure repulsive potential

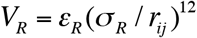

were *r_i,j_* is the inter-bead distance, *σ_R_* = 4 Å, and *ε_R_* = 2.0 *kcal/mol*.

The interaction between quasi-rigid domains is modeled with the residue-specific pair interaction potentials that combine short-range interactions with the long-range electrostatics as it described (27, 28). The short-range interaction is given by a Lennard-Jones 12-10-6-type potential and simple Debye-Hückel-type potential is used for the electrostatics interaction (27). In this study we used the dielectric constant of 80 and the Debye screening length of 10 Å, which corresponds to a salt concentration of about 100 mM.

To account for the experimentally observed cross-links we introduced used in the force field additional distance restraints term given by a potential

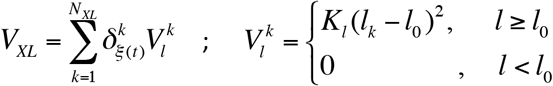

Here sum is over all cross-links, *N_Xl_* is the number of cross-links, *l_k_* is C_α_-C_α_ distance for residues involved in k-th cross-link, *l_0_* = 32 Å is upper bound, *K_XL_* = 10 kcal/mol is force constant, 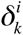 is Kronecker delta, and *ξ(t)* is random digital number selected from the interval [1, *N_Xl_*]. *ξ(t)* is a number that is randomly changed every *τ_XL_* = 10 ns during the MD simulation.

In-house software was developed and used for performing constant temperature molecular dynamics simulations of the coarse-grained model described above. Andersen method (29) was used to control temperature.

### NMR Spectroscopy

All TROSY NMR spectra were collected at 25°C on Bruker Avance(II) 800 MHz spectrometer equipped with ^1^H/^13^C/^15^N cryoprobe. All NMR samples were prepared at pH 7.7 with 20 mM TRIS, 250 mM NaCl, 2mM β-mercaptoethanol, 2 mM DTT, 1 mM PMSF. The final NMR samples contained 5% D_2_O with a protein concentration ranging between 100 μM and 350 μM. The spectra were processed with NMR Pipe software and analyzed with SPARKY (T.D. Goddard and D.G. Kneller, SPARKY 3, University of California, San Francisco). ^1^H-^15^N-ΔN-WDR5 backbone chemical shifts were assigned using ABACUS approach(30) combined with manual analysis using NMR data collected at high resolution from nonlinearly sampled spectra (HNCO, CBCA(CO)NH, HBHA(CO)NH, HNCA, HNCACB), and processed using multidimensional decomposition (31, 32). Figures were prepared using PyMol (DeLano Scientific).

### NMR Titration Experiments

NMR samples were prepared in buffer containing 20 mM TRIS pH 7.4, 150 mM NaCl, 2mM DTT, 1 mM TCEP, 0.5 mM PMSF and 5% D_2_O. Aliquots of MLL1-WIN (GSARAEVHLRKS) and RbBP5 (EDEEVDVTSV) peptides were titrated into the labeled WDR5 protein in molar ratios of 1:1, 1:3, 1:5 and 1:7, until no further changes in chemical shifts were detected in the ^1^H-^15^N TROSY spectra. OICR-9429 added to WDR5 in molar ratios 1:1 and 1:2. The weighted chemical shift perturbations were calculated using following formula: Δppm=[(δ_NH_)^2^+(δ_N_/5)^2^]^1/2^.

### Gel Filtration experiments

A calibrated Superdex 200 column was equilibrated with 20 mM Tris pH 7.7, 150 mM NaCl, 10uM ZnCl_2_, 5mM β-mercaptoethanol, 5 mM DTT, 1 mM phenylmethylsulphonyl fluoride (PMSF). For Fig 6A, WDR5, MLL1-WIN and RbBP5-FL proteins were combined in 2:1:2 molar ratios with protein concentrations 12 μM, 6 μM and 13 μM, respectively and loaded onto the Superdex 200 column (Figs 6A navy blue trace and S3B). For OICR-9429 competition studies (Fig 6A, cyan), the column was also pre-equilibrated with 5-fold excess of OICR9429 and the WDR5/MLL1-WIN/RbBP5-FL complex was pre-incubated with OICR-9429 at 1:5 molar ratio.

### GST Pull-down experiments

Recombinant purified MLL1-GST proteins were incubated with RbBP5 fragments at a 1:2 molar ratio in an assay buffer containing 20mM Tris pH 7.7, 150 mM NaCl, 10μM ZnCl_2_, 5mM β-mercaptoethanol, 5 mM DTT, 1 mM PMSF at 4 °C for 1 hour, followed by incubation with 100 μL of glutathione-Sepharose beads (GE Healthcare) for an additional 1 hour. Protein concentrations were 19-30 μM, 9-19 μM, and 10-15 μM for RbBP5fl, RbBP5-NTD, and MLL wild type/mutant constructs, respectively. The mixture was transferred to a micro-column and extensively washed with assay buffer. Bound proteins were eluted with 30 mM reduced glutathione and detected by SDS-PAGE and Coomassie staining.

### Histone methyltransferase assay

Activity assays were performed in 50 mM Tris-HCl, pH 8.0, 5 mM DTT and 0.01% Triton X-100, using 5 μM ^3^H-SAM and 5 μM Biotin-H3 1-25. Increasing concentrations of RbBP5 were added to 200 nM of MLL1-WDR5 complexes (with either wild-type or mutant MLL1). All reactions were incubated for 90 minutes at room temperature and SPA method was used to determine the activities. Experiments were performed in triplicate. To test the effect of OICR9429 on MLL1 complex, increasing concentration of OICR9429 was incubated with 200 nM MLL1-WDR5 complexes for 20 min before adding 400 nM RbBP5. The activity of the complex was measured as above.

---

**Table S1.** Experimental inter-protein cross-links^a^ collected for trimeric WDR5/RbBP5/MLL1-WIN complex.

| Protein 1 | Protein 2 |
| --- | --- |

| **Cross-links used in the modeling:** |  |
| --- | --- |
| **MLL1<sub>3828</sub>** | **RbBP5<sub>288</sub>** |
| **MLL1<sub>3846</sub>** | **RbBP5<sub>288</sub>** |
| **MLL1<sub>3870</sub>** | **RbBP5<sub>288</sub>** |
| **MLL1<sub>3870</sub>** | **RbBP5<sub>244</sub>** |
| **MLL1<sub>3846</sub>** | **WDR5<sub>46</sub>** |
| **RbBP5<sub>60</sub>** | **WDR5<sub>159</sub>** |
| **Cross-links within flexible regions:** |  |
| **MLL1<sub>3749</sub>** | **WDR5<sub>7</sub>** |
| **MLL1<sub>3749</sub>** | **WDR5<sub>70</sub>** |
| **MLL1<sub>3749</sub>** | **WDR5<sub>46</sub>** |
| **MLL1<sub>3749</sub>** | **RbB5<sub>288</sub>** |
| **MLL1<sub>3749</sub>** | **RbB5<sub>505</sub>** |
| **MLL1<sub>3749</sub>** | **RbB5<sub>517</sub>** |

